# The Alk receptor tyrosine kinase regulates Sparkly, a novel activity regulating neuropeptide precursor in the *Drosophila* CNS

**DOI:** 10.1101/2023.06.02.543395

**Authors:** Sanjay Kumar Sukumar, Vimala Antonydhason, Linnea Molander, Jawdat Sandakly, Malak Kleit, Ganesh Umapathy, Patricia Mendoza-Garcia, Tafheem Masudi, Andreas Schlossser, Dick R. Nässel, Christian Wegener, Margret Shirinian, Ruth H. Palmer

## Abstract

Numerous roles for the Alk receptor tyrosine kinase have been described in *Drosophila*, including functions in the central nervous system (CNS), however the molecular details are poorly understood. To gain mechanistic insight, we employed Targeted DamID (TaDa) transcriptional profiling to identify targets of Alk signaling in the larval CNS. TaDa was employed in larval CNS tissues, while genetically manipulating Alk signaling output. The resulting TaDa data were analysed together with larval CNS scRNA-seq datasets performed under similar conditions, identifying a role for Alk in the transcriptional regulation of neuroendocrine gene expression. Further integration with bulk/scRNA-seq and protein datasets from larval brains in which Alk signaling was manipulated, identified a previously uncharacterized *Drosophila* neuropeptide precursor encoded by *CG4577* as an Alk signaling transcriptional target. *CG4577*, which we named *Sparkly (Spar),* is expressed in a subset of Alk-positive neuroendocrine cells in the developing larval CNS, including circadian clock neurons. In agreement with our TaDa analysis, overexpression of the *Drosophila* Alk ligand Jeb resulted in increased levels of Spar protein in the larval CNS. We show that Spar protein is expressed in circadian (Clock) neurons, and flies lacking Spar exhibit defects in sleep and circadian activity control. In summary, we report a novel activity regulating neuropeptide precursor gene that is regulated by Alk signaling in the *Drosophila* CNS.

## Introduction

Receptor tyrosine kinases (RTK) are involved in wide range of developmental processes. In humans, the Anaplastic Lymphoma Kinase (ALK) RTK is expressed in the central and peripheral nervous system and its role as an oncogene in the childhood cancer neuroblastoma, which arises from the peripheral nervous system, is well described (Iwahara *et al*, 1997; Matthay *et al*, 2016; Umapathy *et al*, 2019; Vernersson *et al*, 2006).

In *Drosophila melanogaster,* Alk is expressed in the visceral mesoderm, central nervous system (CNS) and at neuromuscular junctions (NMJ). The critical role of *Drosophila* Alk and its ligand Jelly belly (Jeb) in the development of the embryonic visceral mesoderm has been extensively studied (Englund *et al*, 2003; Jin *et al*, 2013; Lee *et al*, 2003; Loren *et al*, 2003; Mendoza-Garcia *et al*, 2021; Mendoza-Garcia *et al*, 2017; Pfeifer *et al*, 2022; Popichenko *et al*, 2013; Reim *et al*, 2012; Schaub & Frasch, 2013; Shirinian *et al*, 2007; Stute *et al*, 2004; Varshney & Palmer, 2006; Wolfstetter *et al*, 2017). In the CNS, Alk signaling has been implicated in diverse functions, including targeting of photoreceptor axons in the developing optic lobes (Bazigou *et al*, 2007), regulation of NMJ synaptogenesis and architecture (Rohrbough & Broadie, 2010; Rohrbough *et al*, 2013) and mushroom body neuronal differentiation (Pfeifer *et al*., 2022). In addition, roles for Alk in neuronal regulation of growth and metabolism, organ sparing and proliferation of neuroblast clones, as well as sleep and long-term memory formation in the CNS have been reported (Bai & Sehgal, 2015; Cheng *et al*, 2011; Gouzi *et al*, 2011; Orthofer *et al*, 2020). The molecular mechanisms underlying these Alk-driven phenotypes are currently under investigation, with some molecular components of *Drosophila* Alk signaling in the larval CNS, such as the protein tyrosine phosphatase Corkscrew (Csw), identified in recent BioID-based *in vivo* proximity labeling analyses (Uckun *et al*, 2021).

In this work, we aimed to capture Alk-signaling dependent transcriptional events in the *Drosophila* larval CNS using Targeted DamID (TaDa) that profiles RNA polymerase II (Pol II) occupancy. TaDa employs a prokaryotic DNA adenine methyltransferase (Dam) to specifically methylate adenines within GATC sequences present in the genome, creating unique GA^me^TC marks. In TaDa, expression of Dam fused to Pol II results in GA^me^TC marks on sequences adjacent to the Pol II binding site and can be combined with the Gal4/UAS system to achieve cell-type specific transcriptional profiling (Southall *et al*, 2013). Tissue specific TaDa analysis of Alk signaling, while genetically manipulating Alk signaling output, has previously been used to identify Alk transcriptional targets in the embryonic visceral mesoderm, such as the transcriptional regulator *Kahuli (Kah)* (Mendoza-Garcia *et al*., 2021). Here, we employed this strategy to identify Alk transcriptional targets in *Drosophila* larval brain tissue. These Alk TaDa identified transcripts were enriched in neuroendocrine cells. Further integration with bulk RNA-seq datasets generated from *Alk* gain-of-function and loss-of-function alleles, identified the uncharacterized neuropeptide precursor (*CG4577*), as an Alk target in the *Drosophila* brain, that we have named *Sparkly (Spar)* based on its protein expression pattern. Spar is expressed in a subset of Alk-expressing cells in the central brain and ventral nerve cord, overlapping with the expression pattern of neuroendocrine specific transcription factor Dimmed (Dimm) (Hewes *et al*, 2003). Further, using genetic manipulation of Alk we show that Spar levels in the CNS respond to Alk signaling output, validating *Spar* as a transcriptional target of Alk. *Spar* mutant flies showed significant reduction in life-span, and behavioral phenotypes including defects in activity, sleep, and circadian activity. Notably, *Alk* loss-of-function alleles displayed similar behavioral defects, suggesting that Alk-dependant regulation of Spar in peptidergic neuroendocrine cells modulates activity and sleep/rest behavior. Interestingly, Alk and its ligand Alkal2 play a role in regulation of behavioral and neuroendocrine function in vertebrates (Ahmed *et al*, 2022; Bilsland *et al*, 2008; Borenas *et al*, 2021; Lasek *et al*, 2011a; Lasek *et al*, 2011b; Orthofer *et al*., 2020; Weiss *et al*, 2012; Witek *et al*, 2015). Taken together, our findings suggest an evolutionarily conserved role of Alk signaling in the regulation of neuroendocrine cell function and identify *Spar* as the first molecular target of Alk to be described in the regulation of activity and circadian control in the fly.

## Results

### TaDa identifies Alk-regulated genes in *Drosophila* larval CNS

To characterize Alk transcriptional targets in the *Drosophila* CNS we employed Targeted DamID (TaDa). Briefly, transgenic DNA adenine methyltransferase (Dam) fused with RNA-Pol II (here after referred as Dam-Pol II) (Southall *et al*., 2013) **(Figure 1a-b)**, was driven using the pan neuronal *C155-Gal4* driver. To inhibit Alk signaling we employed a dominant negative Alk transgene, which encodes the Alk extracellular and transmembrane domain (here after referred as *UAS-Alk^DN^*) (Bazigou *et al*., 2007) **(Figure 1a)**. Flies expressing Dam-Pol II alone in a wild-type background were used as control. Expression of Dam-Pol II was confirmed by expression of mCherry, which is encoded by the primary ORF of the TaDa construct (Southall *et al*., 2013) **(Figure 1b, Figure 1 – figure supplement 1a-b’)**. CNS from third instar wandering larvae were dissected and genomic DNA was extracted, fragmented at GA^me^TC marked sites using methylation specific DpnI restriction endonuclease. The resulting GATC fragments were subsequently amplified for library preparation and NGS sequencing **(Figure 1 – figure supplement 1c)**. Bioinformatic data analysis was performed based on a previously described pipeline (Marshall & Brand, 2017; Mendoza-Garcia *et al*., 2021). Initial quality control analysis indicated comparable numbers of quality reads between samples and replicates, identifying >20 million raw reads per sample that aligned to the *Drosophila* genome **(Figure 1 – figure supplement 1d-d’)**. No significant inter-replicate variability was observed **(Figure 1 – figure supplement 1e)**. Meta-analysis of reads associated with GATC borders showed a tendency to accumulate close to Transcription Start Sites (TSS) indicating the ability of TaDa to detect transcriptionally active regions **(Figure 1 – figure supplement 1f)**. A closer look at the Pol II occupancy profile of *Alk* shows a clear increase in Pol II occupancy from Exon 1 to Exon 7 (encoding the extracellular and transmembrane domain) in *Alk^DN^* samples reflecting the expression of the dominant negative *Alk* transgene **(Figure 1 – figure supplement 1g)**.

**Figure 1.**
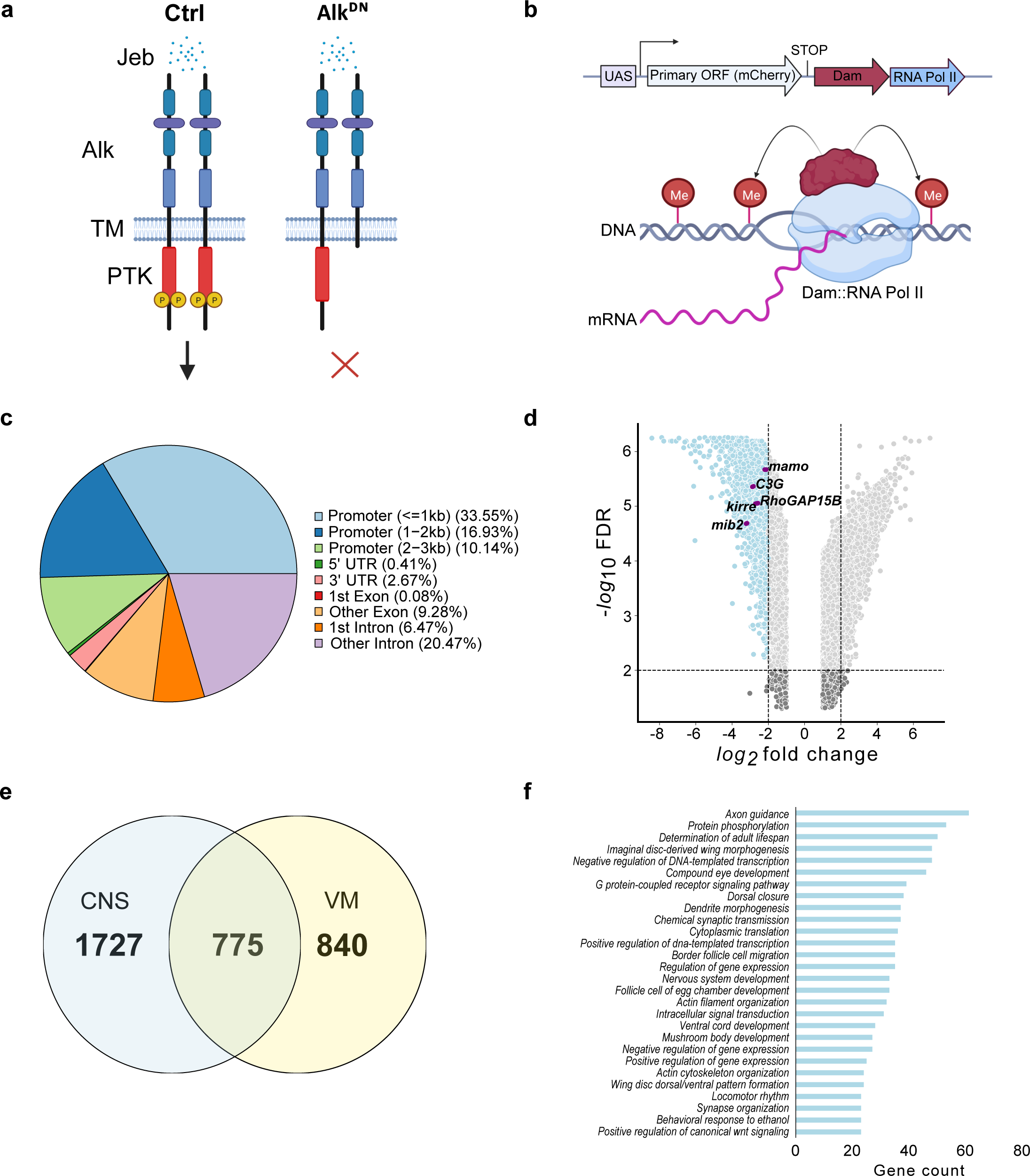
TaDa-seq identifies novel Alk-regulated genes in the *Drosophila* larval CNS. **a.** Schematic overview of experimental conditions comparing wild-type Alk (Ctrl) with Alk dominant-negative (*Alk^DN^)* conditions. The *Drosophila* Alk RTK is comprised of extracellular, transmembrane and intracellular kinase (red) domains. Upon Jelly belly (Jeb, blue dots) ligand stimulation the Alk kinase domain is auto-phosphorylated (yellow circles) and downstream signaling is initiated. In *Alk^DN^* experimental conditions, Alk signaling is inhibited due to overexpression of the Alk extracellular domain. **b.** The TaDa system (expressing *Dam::RNA Pol II*) leads to the methylation of GATC sites in the genome, allowing transcriptional profiling based on RNA Pol II occupancy. **c**. Pie chart indicating the distribution of TaDa peaks on various genomic features such as promoters, 5’ UTRs, 3’ UTRs, exons and introns. **d.** Volcano plot of TaDa-positive loci enriched in *Alk^DN^* experimental conditions compared to control loci exhibiting Log2FC<-2, *p*≥0.05 are shown in blue. Alk-associated genes such as *mamo, C3G, Kirre, RhoGAP15B* and *mib2* are highlighted in purple. **e.** Venn diagram indicating Alk-dependant TaDa downregulated genes from the current study compared with previously identified Alk-dependant TaDa loci in the embryonic VM (Mendoza-Garcia *et al*., 2021). **f.** Enrichment of Gene Ontology (GO) terms associated with significantly down-regulated genes in *Alk^DN^* experimental conditions.

To detect differential Pol II occupancy between Dam-Pol II control (*C155-Gal4>UAS-LT3-Dam::Pol ll*) and *UAS-Alk^DN^* (*C155-Gal4>UAS-LT3-Dam::Pol II; UAS-Alk^DN^*) samples, neighbouring GATC associated reads, maximum 350 bp apart (median GATC fragment distance in the *Drosophila* genome) were clustered in peaks (Tosti *et al*, 2018). More than 10 million reads in both control and *Alk^DN^* samples were identified as GATC associated reads (**Figure 1 – figure supplement 1d’)**, and those loci displaying differential Pol II occupancy were defined by logFC and FDR (as detailed in materials and methods). Greater than 50% of aligned reads were in promoter regions, with 33.55% within a 1 kb range **(Figure 1c, Table S1)**.

To further analyse transcriptional targets of Alk signaling we focused on loci exhibiting decreased Pol II occupancy when compared with controls, identifying 2502 loci with logFC<-2, FDR<0.05 **(Figure 1d, Table S1)**. Genes previously known to be associated with Alk signaling, such as *kirre, RhoGAP15B, C3G, mib2* and *mamo*, were identified among downregulated loci **(Figure 1d)**. We compared CNS TaDa Alk targets with our previously published embryonic visceral mesoderm TaDa datasets that were derived under similar experimental conditions (Mendoza-Garcia *et al*., 2021) and found 775 common genes **(Figure 1e, Table S1)**. Gene ontology (GO) analysis identified GO terms in agreement with previously reported Alk functions in the CNS (Bai & Sehgal, 2015; Bazigou *et al*., 2007; Cheng *et al*., 2011; Gouzi *et al*., 2011; Orthofer *et al*., 2020; Pfeifer *et al*., 2022; Rohrbough & Broadie, 2010; Rohrbough *et al*., 2013; Woodling *et al*, 2020) such as axon guidance, determination of adult lifespan, nervous system development, regulation of gene expression, mushroom body development, behavioral response to ethanol and locomotor rhythm (**Figure 1f**). Many of the differentially regulated identified loci have not previously been associated with Alk signaling and represent candidates for future characterisation.

### TaDa targets are enriched for neuroendocrine transcripts

To further characterize Alk-regulated TaDa loci, we set out to examine their expression in scRNA-seq data from wild-type third instar larval CNS (Pfeifer *et al*., 2022). Enrichment of TaDa loci were identified by using AUCell, an area-under-the-curve based enrichment score method, employing the top 500 TaDa hits (Aibar *et al*, 2017) **(Figure 2a-b, Table S1)**. This analysis identified 786 cells (out of 3598), mainly located in a distinct cluster of mature neurons that robustly express both *Alk* and *jeb* **(Figure 2b, red circle; Figure 2c)**. This cluster was defined as neuroendocrine cells based on canonical markers, such as the neuropeptides *Lk* (*Leucokinin*), *Nplp1* (*Neuropeptide-like precursor 1*), *Dh44* (*Diuretic hormone 44*), *Dh31* (*Diuretic hormone 31*), *sNPF* (*short neuropeptide F*), *AstA* (*Allatostatin A*), and the enzyme *Pal2* (*Peptidyl-α-hydroxyglycine-α-amidating lyase 2*) as well as *Eip74EF* (*Ecdysone-induced protein 74EF*), and *Rdl* (resistance to dieldrin) (Guo *et al*, 2019; Huckesfeld *et al*, 2021; Takeda & Suzuki, 2022; Torii, 2009) **(Figure 2d-f)**. Overall, the TaDa-scRNAseq data integration analysis suggests a role of Alk signaling in regulation of gene expression in neuroendocrine cells.

**Figure 2.**
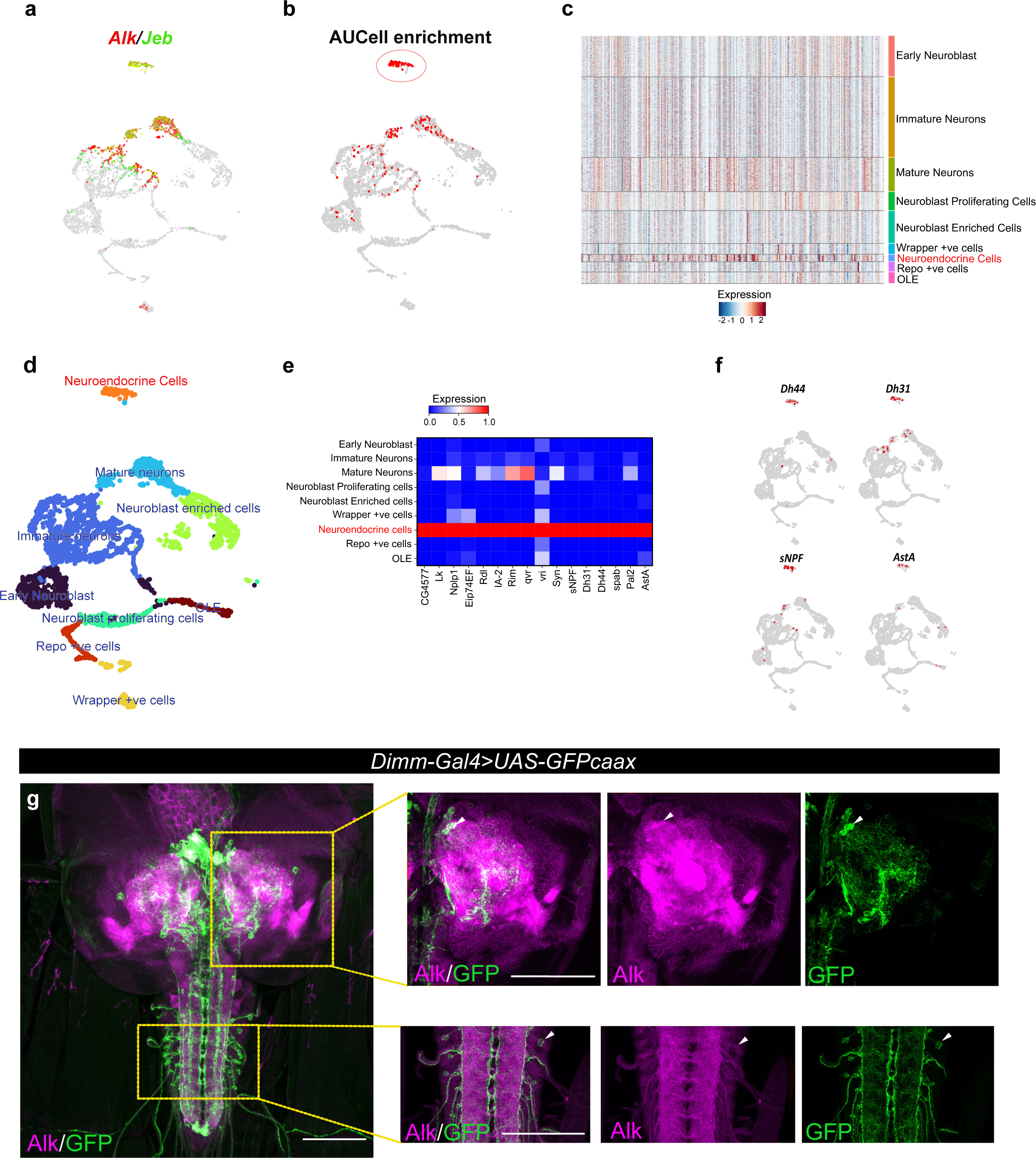
Integration of TaDa data with scRNA-seq identifies an enrichment of Alk-regulated genes in neuroendocrine cells. **a.** UMAP feature plot indicating Alk (in red) and Jeb (in green) mRNA expression in a control (*w^1118^*) whole third instar larval CNS scRNA-seq dataset (Pfeifer *et al*., 2022). **b.** UMAP visualizing AUCell enrichment analysis of the top 500 TaDa downregulated genes in the third instar larval CNS scRNA-seq dataset. Cells exhibiting an enrichment (threshold >0.196) are depicted in red. One highly enriched cell cluster is highlighted (red circle). **c.** Heatmap representing expression of the top 500 genes downregulated in TaDa *Alk^DN^* samples across larval CNS scRNA-seq clusters identifies enrichment in neuroendocrine cells. **d.** UMAP indicating third instar larval CNS annotated clusters (Pfeifer *et al*., 2022), including the annotated neuroendocrine cell cluster (in orange). **e.** Matrix plot displaying expression of canonical neuroendocrine cell markers. **f.** Feature plot visualizing mRNA expression of *Dh44*, *Dh31, sNPF* and *AstA* neuropeptides across the scRNA population. **g.** Alk staining in *Dimm-Gal4>UAS-GFPcaax* third instar larval CNS confirms Alk expression in Dimm-positive cells. Alk (in magenta) and GFP (in green), close-ups indicated by boxed regions and arrows indicating overlapping cells in the central brain and ventral nerve cord. Scale bars: 100 μm.

To further explore the observed enrichment of Alk-regulated TaDa loci in neuroendocrine cells, we used a Dimmed (Dimm) transcription factor reporter (*Dimm-Gal4>UAS-GFPcaax*), as a neuroendocrine marker (Park *et al*, 2008), to confirm Alk protein expression in a subset of neuroendocrine cells in the larval central brain, ventral nerve cord and neuroendocrine corpora cardiaca cells **(Figure 2g, Figure 2 – figure supplement 1)**. This could not be confirmed at the RNA level, due to low expression of *dimm* in both our and publicly available single cell RNASeq datasets (Brunet Avalos *et al*, 2019; Michki *et al*, 2021; Pfeifer *et al*., 2022).

### Multi-omics integration identifies *CG4577* as an Alk transcriptional target

Loci potentially subject to Alk-dependent transcriptional regulation were further refined by integration of the Alk-regulated TaDa dataset with previously collected RNA-seq datasets **(Figure 3a)**. Specifically, *w^1118^* (control), *Alk^Y1355S^* (Alk gain-of-function) and *Alk^ΔRA^* (Alk loss-of-function) RNA-seq datasets (Pfeifer *et al*., 2022) were compared to identify genes that exhibited both significantly increased expression in Alk gain-of-function conditions (*w^1118^* vs *Alk^Y1355S^*) and significantly decreased expression in Alk loss-of-function conditions (*w^1118^* vs *Alk^ΔRA^* and control vs *C155-Gal4* driven expression *of UAS-Alk^DN^*). Finally, we positively selected for candidates expressed in *Alk*-positive cells in our scRNA-seq dataset. Notably, the only candidate which met these stringent criteria was *CG4577*, which encodes an uncharacterized putative neuropeptide precursor (**Figure 3b)**. *CG4577* exhibited decreased Pol II occupancy in *Alk^DN^* samples **(Figure 3c),** and *CG4577* transcripts were upregulated in *Alk^Y1355S^* gain-of-function conditions and downregulated in *Alk^ΔRA^* loss-of-function conditions (**Figure 3d, Table S1).** In agreement with a potential role as a neuropeptide precursor, expression of *CG4577* was almost exclusively restricted to neuroendocrine cell clusters in our scRNA-seq dataset **(Figure 3e)**. Examination of additional publicly available first instar larval and adult CNS scRNAseq datasets (Brunet Avalos *et al*., 2019; Davie *et al*, 2018) confirmed the expression of *CG4577* in Alk-expressing cells **(Figure 3 – figure supplement 1a-b)**. *CG4577-RA* encodes a 445 amino acid prepropeptide with a 27 aa N-terminal signal peptide sequence as predicted by SignalP-5.0 **(Figure 3f)** (Almagro Armenteros *et al*, 2019). Analysis of CG4577-PA at the amino acid level identified a high percentage of glutamine residues (43 of 445; 9%), including six tandem glutamine repeats (amino acids 48-56, 59-62, 64-71, 116-118, 120-122 and 148-150) of unknown function as well as a lack of cysteine residues. The preproprotein has an acidic pI of 5.1 and carries a net negative charge of 6. Several poly- and di-basic prohormone convertase (PC) cleavage sites were also predicted (KR, KK, RR, RK) (Pauls *et al*, 2014; Southey *et al*, 2006; Veenstra, 2000) **(Figure 3f)**. Since the propeptide does not contain cysteine residues it is unable to form intracellular or dimeric disulfide bridges. A second transcript, *CG4577-RB*, encodes a 446 amino acid protein with only two amino acid changes **(Figure 3 – figure supplement 1c).** Phylogenetic analysis of *CG4577* relative to known *Drosophila* neuropeptide precursors failed to identify strong homology in keeping with the known low sequence conservation of neuropeptide prepropeptides outside the bioactive peptide stretches. However, we were also unable to find sequence homologies with other known invertebrate or vertebrate peptides. Next, we searched for *CG4577* orthologs across Metazoa. We obtained orthologs across the Drosophilids, Brachyceran flies and Dipterans. No orthologs were found at higher taxonomic levels, suggesting that *CG4577* either originated in Dipterans, or has a high sequence variability at higher taxonomic levels. To identify conserved peptide stretches indicating putative bioactive peptide sequences, we aligned the predicted aa sequences of the Dipteran *CG4577* orthologs. This revealed several conserved peptide stretches (**Figure 3 – figure supplement 2**) framed by canonical prohormone cleavage sites that might represent bioactive peptide sequences. BLAST searches against these conserved sequences did not yield hits outside of the Diptera.

**Figure 3.**
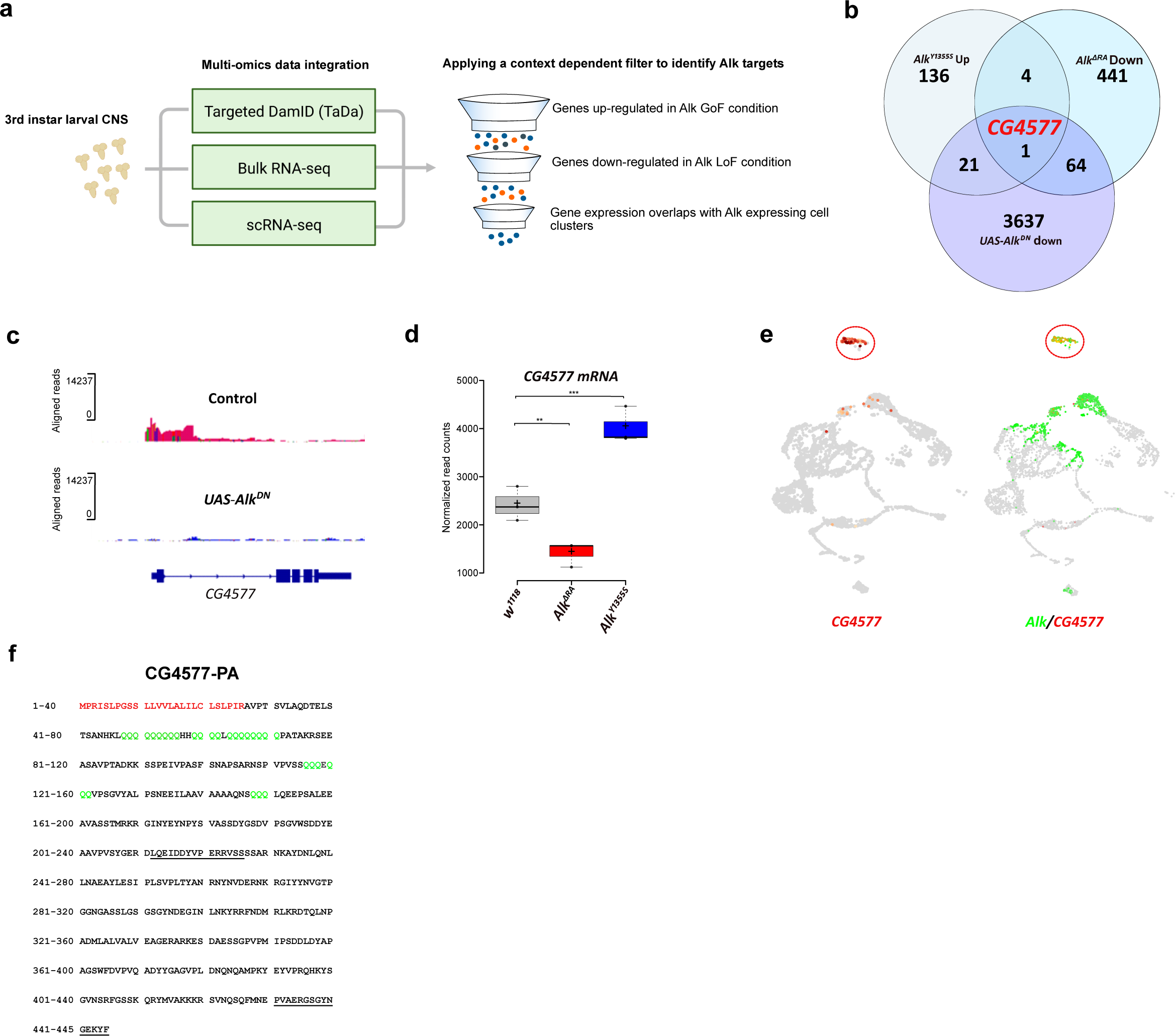
TaDa and RNA-seq identifies CG4577 as a novel Alk-regulated neuropeptide. **a.** Flowchart representation of the multi-omics approach employed in the study and the context dependent filter used to integrate TaDa, bulk RNA-seq and scRNA-seq datasets. **b.** Venn diagram comparing bulk RNA-seq (Log2FC>1.5, *p*<0.05) and TaDa datasets (Log2FC<-2, *p*<0.05). A single candidate (*CG4577/Spar*) is identified as responsive to Alk signaling. **c.** TaDa Pol II occupancy of *CG4577/Spar* shows decreased occupancy in *Alk^DN^* experimental conditions compared to control. **d.** Expression of *CG4577/Spar* in *w^1118^* (control), *Alk^ΔRA^* (*Alk* loss-of-function allele) and *Alk^Y1355S^* (*Alk* gain-of-function allele) larval CNS. Boxplot with normalized counts, \*\**p*<0.01, \*\*\**p*<0.001. **e.** Feature plot showing mRNA expression of *CG4577/Spar* and *Alk* in third instar larval CNS scRNA-seq data. Neuroendocrine cluster is highlighted (red circle). **f.** CG4577/Spar-PA amino acid sequence indicating the signal peptide (amino acids 1-26, in red), glutamine repeats (in green) and the anti-CG4577/Spar antibody epitopes (amino acids 211-225 and 430-445, underlined). Center lines in boxplots indicate medians; box limits indicate the 25th and 75th percentiles; crosses represent sample means; whiskers extend to the maximum or minimum.

### CG4577/Spar is expressed in neuroendocrine cells

To further characterize *CG4577* we generated polyclonal antibodies that are predicted to recognize both CG4577-PA and CG4577-PB and investigated protein expression. CG4577 protein was expressed in a “sparkly” pattern in neurons of the third instar central brain as well as in distinct cell bodies and neuronal processes in the ventral nerve cord, prompting us to name CG4577 as Sparkly (Spar) (**Figure 4a-b**). Co-labeling of Spar and Alk confirmed expression of Spar in a subset of Alk-expressing cells, in agreement with our transcriptomics analyses **(Figure 4a, Figure 4 – figure supplement 1)**. In addition, we also observed expression of Spar in neuronal processes which emerge from the ventral nerve cord and appear to innervate larval body wall muscle number 8, that may be either Leukokinin (Lk) or cystine-knot glycoprotein hormone GPB5 expressing neurons **(Figure 4b)** (Cantera & Nässel, 1992; Sellami *et al*, 2011). Spar antibody specificity was confirmed in both *C155-Gal4>UAS-Spar-RNAi* larvae, where RNAi-mediated knock down of *Spar* resulted in loss of detectable signal (**Figure 4c-c’**), and in *C155-Gal4>UAS-Spar* larvae, exhibiting ectopic *Spar* expression in the larval CNS and photoreceptors of the eye disc (**Figure 4d-d’**). To further address Spar expression in the neuroendocrine system, we co-labelled with antibodies against Dimm to identify peptidergic neuronal somata (Allan et al., 2005) in a *Dimm-Gal4>UAS-GFPcaax* background. This further confirmed the expression of Spar in Dimm-positive peptidergic neuroendocrine cells in the larval CNS (**Figure 4e-e’’, Figure 4 – Movie supplements 1 and 2**). Moreover, co-staining of Spar and Dimm in the adult CNS showed similar results **(Figure 4 – figure supplement 2)**.

**Figure 4.**
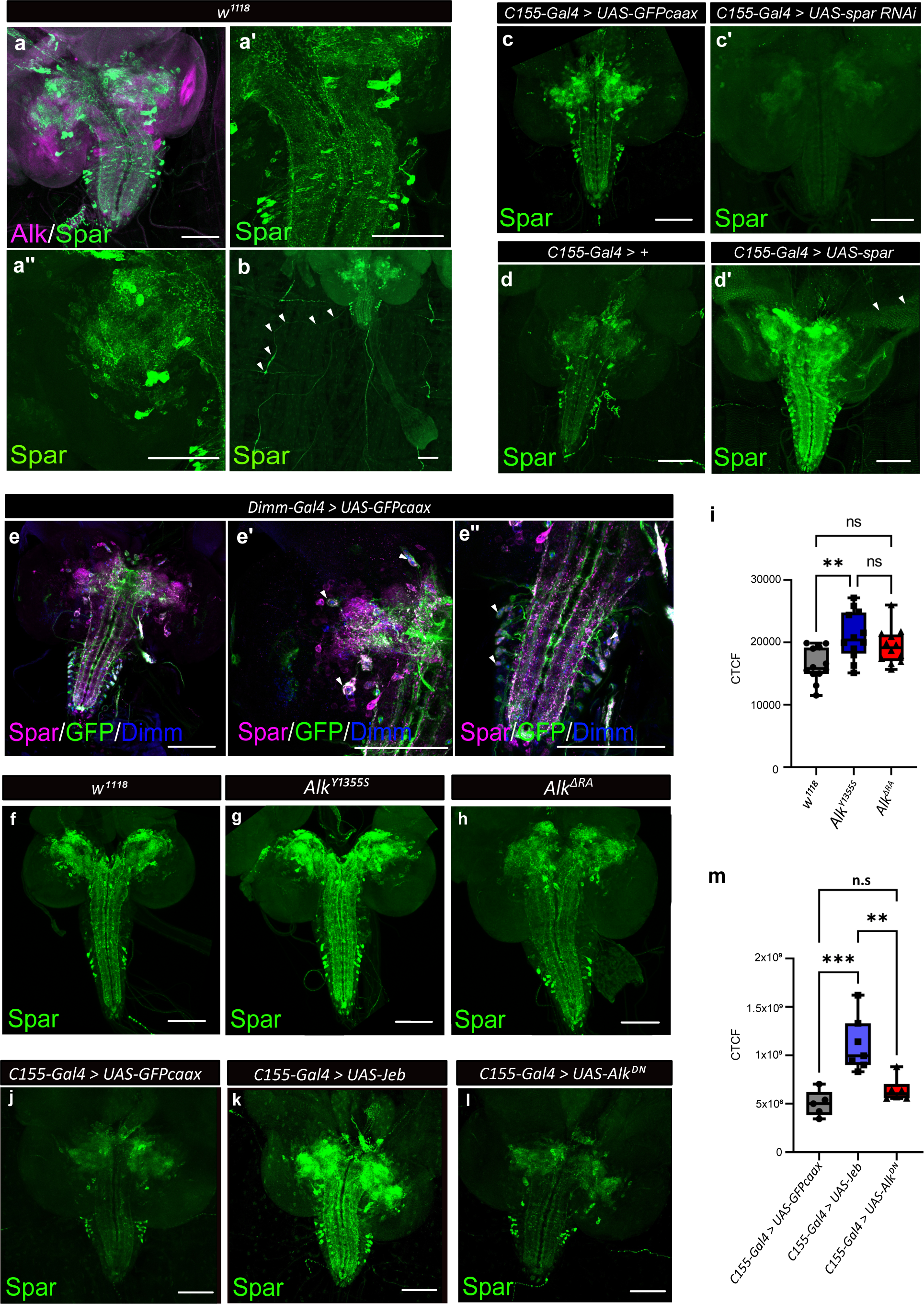
Spar expression in the Drosophila larval brain. **a**. Immunostaining of *w^1118^* third instar larval brains with Spar (green) and Alk (magenta) revealing overlapping expression in central brain and ventral nerve cord. **a’-a’’**. Close-up of Spar expression (green) in ventral nerve cord (a’) and central brain (a’’). **b**. Immunostaining of *w^1118^* third instar larval CNS together with the body wall muscles, showing Spar (green) expression in neuronal processes (white arrowheads) which emerge from the ventral nerve cord and innervate larval body wall muscle number 8. **c-c’**. Decreased expression of Spar in third instar larval brains expressing *spar* RNAi (*C155-Gal4>Spar RNAi*) compared to control (*C155-Gal4>UAS-GFPcaax*) confirms Spar antibody specificity (Spar in green). **d-d’**. Spar overexpression (*C155-Gal4>UAS-Spar*) showing increased Spar expression (in green) compared to control (*C155-Gal4>+*) larval CNS. **e-e’’**. Immunostaining of *Dimm-Gal4>UAS-GFPcaax* third instar larval brains with Spar (in magenta), GFP and Dimm (in blue) confirms Spar expression in Dimm-positive neuroendocrine cells (white arrowheads). **f-i**. Spar protein expression in *w^1118^*, *Alk^Y1355S^*, and *Alk^ΔRA^* third instar larval brains. Quantification of Spar levels (corrected total cell fluorescence, CTCF) in **i**. **j-m**. Overexpression of Jeb in the third instar CNS (*C155-Gal4>UAS-Jeb*) leads to increased Spar protein expression compared to controls (*C155-Gal4>UAS-GFPcaax*). Quantification of Spar levels (corrected total cell fluorescence, CTCF) in **m**. (\*\**p*<0.01; \*\*\**p*<0.001) Scale bars: 100 μm. Center lines in boxplots indicate medians; box limits indicate the 25th and 75th percentiles; whiskers extend to the maximum or minimum.

### Spar expression is modulated in response to Alk signaling activity

Our initial integrated analysis predicted *Spar* as a locus responsive to Alk signaling. To test this hypothesis, we examined Spar protein expression in *w^1118^*, *Alk^Y1355S^* and *Alk^ΔRA^* genetic backgrounds, in which Alk signaling output is either upregulated (*Alk^Y1355S^*) or downregulated (*Alk^ΔRA^*) (Pfeifer *et al*., 2022). We observed a significant increase in Spar protein in *Alk^Y1355S^* CNS, while levels of Spar in *Alk^ΔRA^* CNS were not significantly altered (**Figure 4f-h**, quantified in **I**). In agreement, overexpression of Jeb *(C155-Gal4>jeb)* significantly increased Spar levels when compared with controls *(C155-Gal4>UAS-GFPcaax)* (**Figure 4j-l**, quantified in **m)**. Again, overexpression of dominant-negative Alk *(C155-Gal4>UAS-Alk^DN^)* did not result in significantly decreased Spar levels (**Figure 4l**, quantified in **m**). Thus activation of Alk signaling increases Spar protein levels. However, while our bulk RNA-seq and TaDa datasets show a reduction in *Spar* transcript levels in Alk loss-of-function conditions, this reduction is not reflected at the protein level. This observation may reflect additional uncharacterized pathways that regulate *Spar* mRNA levels as well as translation and protein stability, since and notably *Spar* transcript levels are decreased but not absent in *Alk^ΔRA^* **(Figure 3d)**. Taken together, these observations confirm that *Spar* expression is responsive to Alk signaling in CNS, although Alk is not critically required to maintain Spar protein levels.

### *Spar* encodes a canonically processed neurosecretory protein

To provide biochemical evidence for the expression of Spar, we re-analysed data from a previous LC-MS peptidomic analysis of brain extracts from five day old male control flies and flies deficient for carboxypeptidase D (dCPD, SILVER) (Pauls *et al*, 2019), an enzyme that removes the basic C-terminal aa of peptides originating from proprotein convertases (PCs) cleavage of the proprotein. This analysis identified several peptides derived from the Spar propeptide by mass matching in non-digested extracts from genetic control brains (**Figure 5**). These included peptides that are framed by dibasic prohormone cleavage sequences in the propeptide, one of which (SEEASAVPTAD) was also obtained by *de-novo* sequencing (**Figure 5**). This result demonstrates that the Spar precursor is expressed and is processed into multiple peptides by PCs and possibly also other proteases. Analysis of the brain of *svr* mutant flies yielded similar results, but further revealed peptides C-terminally extended by the dibasic cleavage sequence (SEEASAVPTADKK, FNDMRLKR) (**Figure 5**), thereby confirming canonical PC processing of the Spar propeptide. Of note, the phylogenetically most conserved peptide sequence of the Spar precursor (DTQLNPADMLALVALVEAGERA, **Figure 3 - figure supplement 2**) framed by dibasic cleavage sites was among the identified peptides yet occurred only in control but not *svr* mutant brains (**Figure 5**).

**Figure 5.**
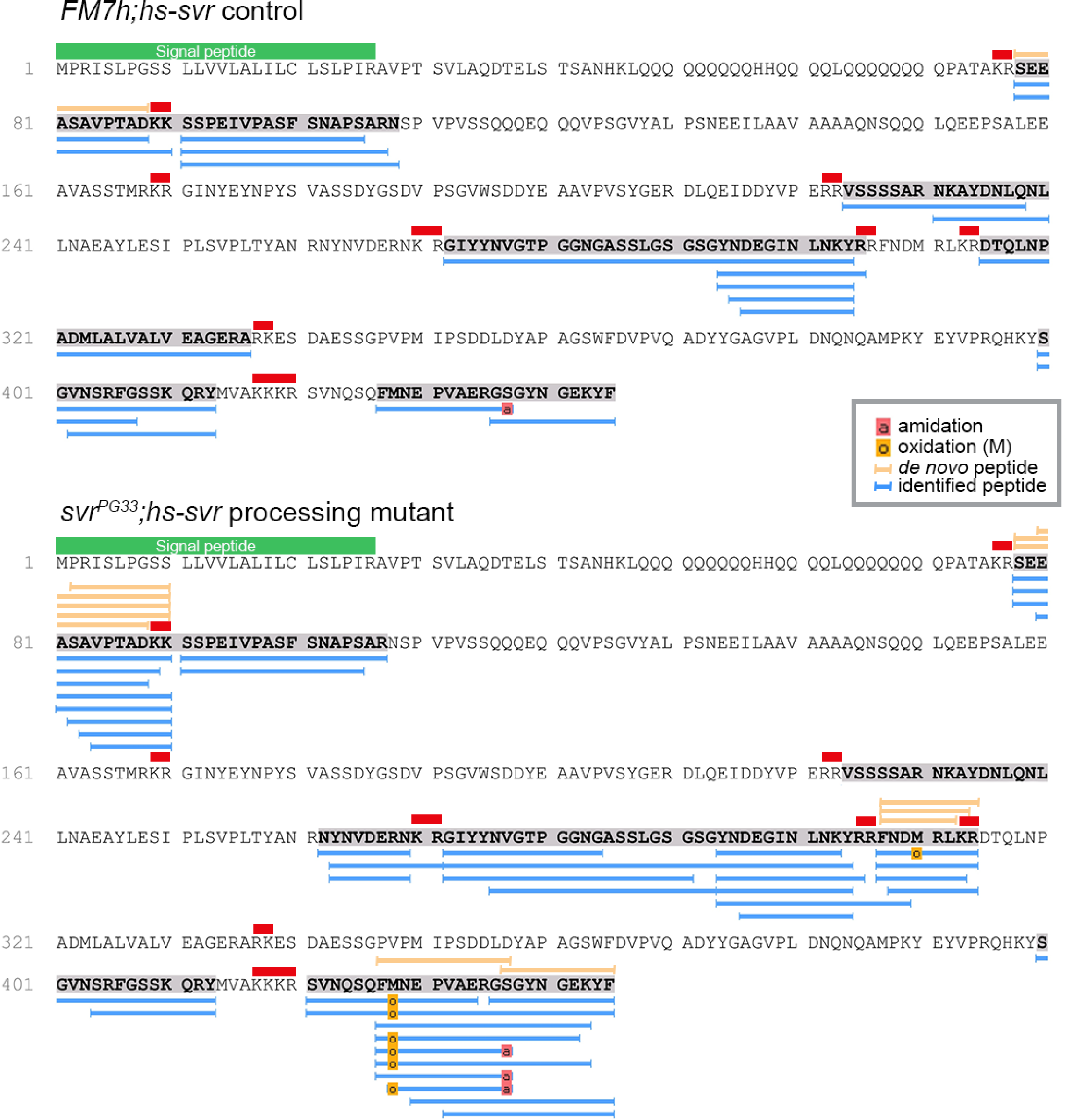
Identification of Spar peptides in *Drosophila* CNS tissues. Peptides derived from the Spar prepropeptide identified by mass spectrometry in wild-type-like control flies (*FM7h;hs-svr*, upper panel) and *svr* mutant (*svrPG33;hs-svr*, lower panel) flies. The predicted amino acid sequence of the CG4577-PA Spar isoform is depicted for each genetic experimental background. Peptides identified by database searching (UniProt *Drosophila melanogaster*, 1% FDR) are marked by blue bars below the sequence. In addition, peptides correctly identified by *de novo* sequencing are marked by orange bars above the sequence. Red bars indicate basic prohormone convertase cleavage sites, green bar indicates the signal peptide.

Additionally, we performed co-labeling with known *Drosophila* neuropeptides, Pigment-dispersing factor (PDF), Dh44, Insulin-like peptide 2 (Ilp2), AstA and Lk, observing Spar expression in subsets of all these populations (**Figure 6**). These included the PDF-positive LNv clock neurons (**Figure 6a-b’’**), Dh44-positive neurons (**Figure 6c-d’’**), a subset of Ilp2 neurons in the central brain (**Figure 6e-f’’**) and several AstA-positive neurons in the central brain and ventral nerve cord (**Figure 6g-h’’**). We also noted co-expression in some Lk-positive neurons in the central brain and ventral nerve cord, that include the neuronal processes converging on body wall muscle 8 (**Figure 6i-l’’**) (Cantera & Nässel, 1992). Similar Spar co-expression with PDF, Dh44, Ilp2, and AstA was observed in adult CNS (**Figure 6 – figure supplement 1**).

**Figure 6.**
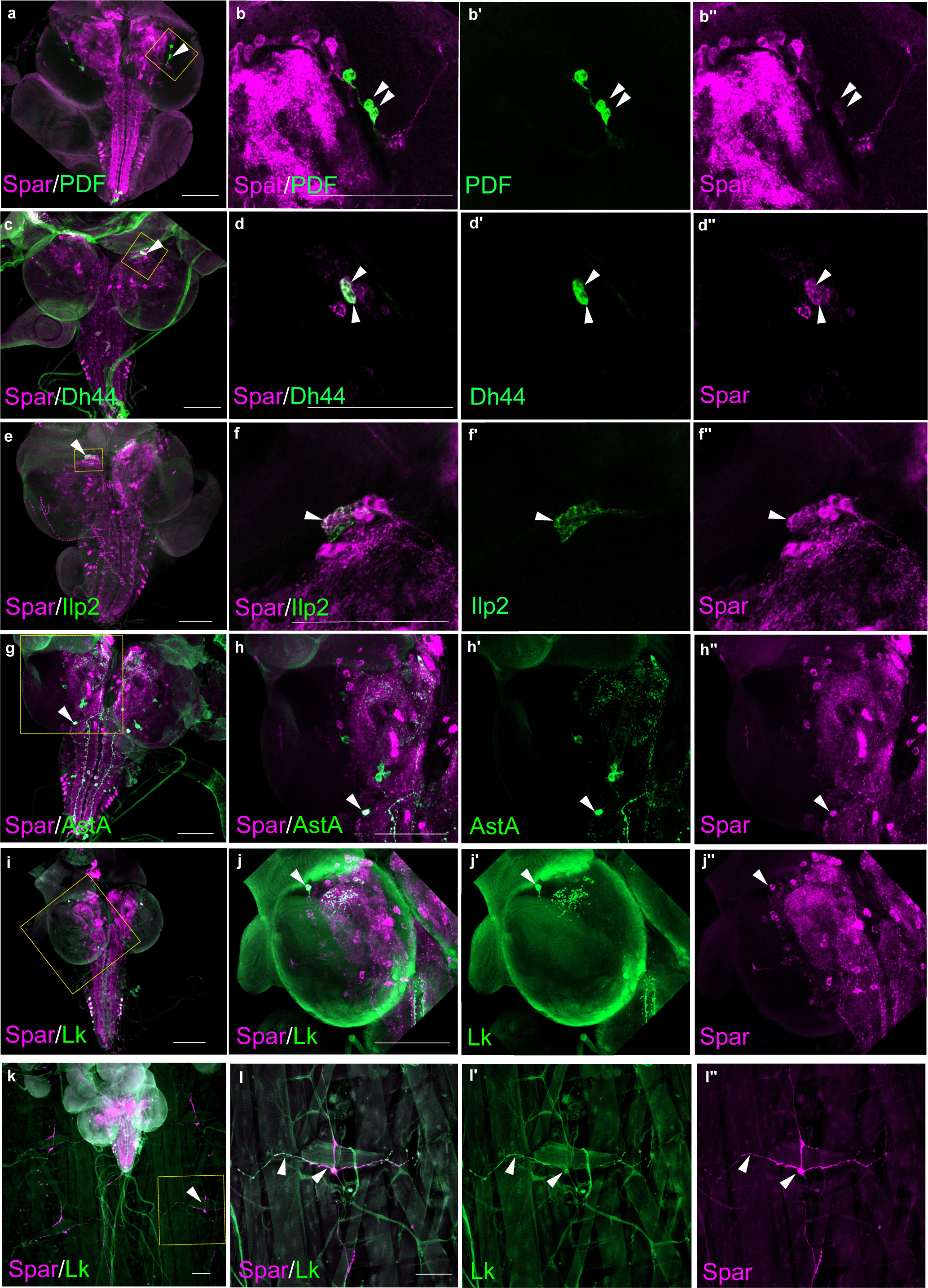
*Spar* expression in larval neuropeptide expressing neuronal populations. **a.** Immunostaining of *w^1118^* third instar larval CNS with Spar (in magenta) and PDF (in green). Closeups (**b-b**’’) showing PDF- and Spar-positive neurons in central brain indicated by white arrowheads. **c**. Immunostaining of *w^1118^* third instar larval CNS with Spar (in magenta) and Dh44 (in green). Closeups (**d-d**’’) showing Dh44- and Spar-positive neurons in central brain indicated by white arrowheads. e. Immunostaining of *w^1118^* third instar larval CNS with Spar (in magenta) and Ilp2 (in green). Closeups (**f-f**’’) showing Ilp2- and Spar-positive neurons in central brain indicated by white arrowheads. **g**. Immunostaining of *w^1118^* third instar larval CNS with Spar (in magenta) and AstA (in green). Closeups (**h-h**’’) showing AstA- and Spar-positive neurons in central brain indicated by white arrowheads. **i**. Immunostaining of *w^1118^* third instar larval CNS with Spar (in magenta) and Lk (in green). Closeups (**j-j**’’) showing Lk (LHLK neurons)- and Spar-positive neurons in central brain indicated by white arrowheads. **k**. Immunostaining of *w^1118^* third instar larval CNS together with the body wall muscles, showing Spar (in magenta) expressing Lk (in green) (ABLK neurons) in neuronal processes, which emerge from the ventral nerve cord and innervate the larval body wall muscle. Closeups (**l-l**’’) showing co-expression of Lk and Spar in neurons which attach to the body wall number 8 indicated by white arrow heads. Scale bars: 100 μm.

### CRISPR/Cas9 generated *Spar* mutants are viable

Since previous reports have shown that Jeb overexpression in the larval CNS results in a small pupal size (Gouzi *et al*., 2011), we measured pupal size on ectopic expression of Spar (*C155-Gal4>Spar*) and *Spar RNAi* (*C155-Gal4>Spar RNAi*), noting no significant difference compared to controls (*C155-Gal4>+* and *C155-Gal4>jeb*) **(Figure 7 – figure supplement 1)**. These results suggest that Spar may be involved in an additional Alk-dependant function in the CNS. Further, experiments overexpressing Spar did not reveal any obvious phenotypes. To further investigate the function of Spar we generated a *Spar* loss of function allele by CRISPR/Cas9-mediated non-homologous end-joining, resulting in the deletion of a 716bp region including the *Spar* transcription start site and exon 1 (hereafter referred as *Spar^ΔExon1^*) (**Figure 7a**). Immunoblotting analysis indicated a 35kDa protein present in the wild-type (*w^1118^*) controls that was absent in *Spar^ΔExon1^* mutant CNS lysates (**Figure 7b**). The *Spar^ΔExon1^* mutant allele was further characterized using immunohistochemistry (**Figure 7c-d’**). *Spar^ΔExon1^* shows a complete abrogation of larval and adult Spar expression, consistent with the reduction observed when *Spar* RNAi was employed (**Figure 7c-d’**). *Spar^ΔExon1^* flies were viable, and no gross morphological phenotypes were observed, similar to loss of function mutants in several previously characterized neuropeptides such as Pigment-dispersing factor (PDF), Drosulfakinin (Dsk) and Neuropeptide F (NPF) (Liu *et al*, 2019; Renn *et al*, 1999; Wu *et al*, 2020).

**Figure 7.**
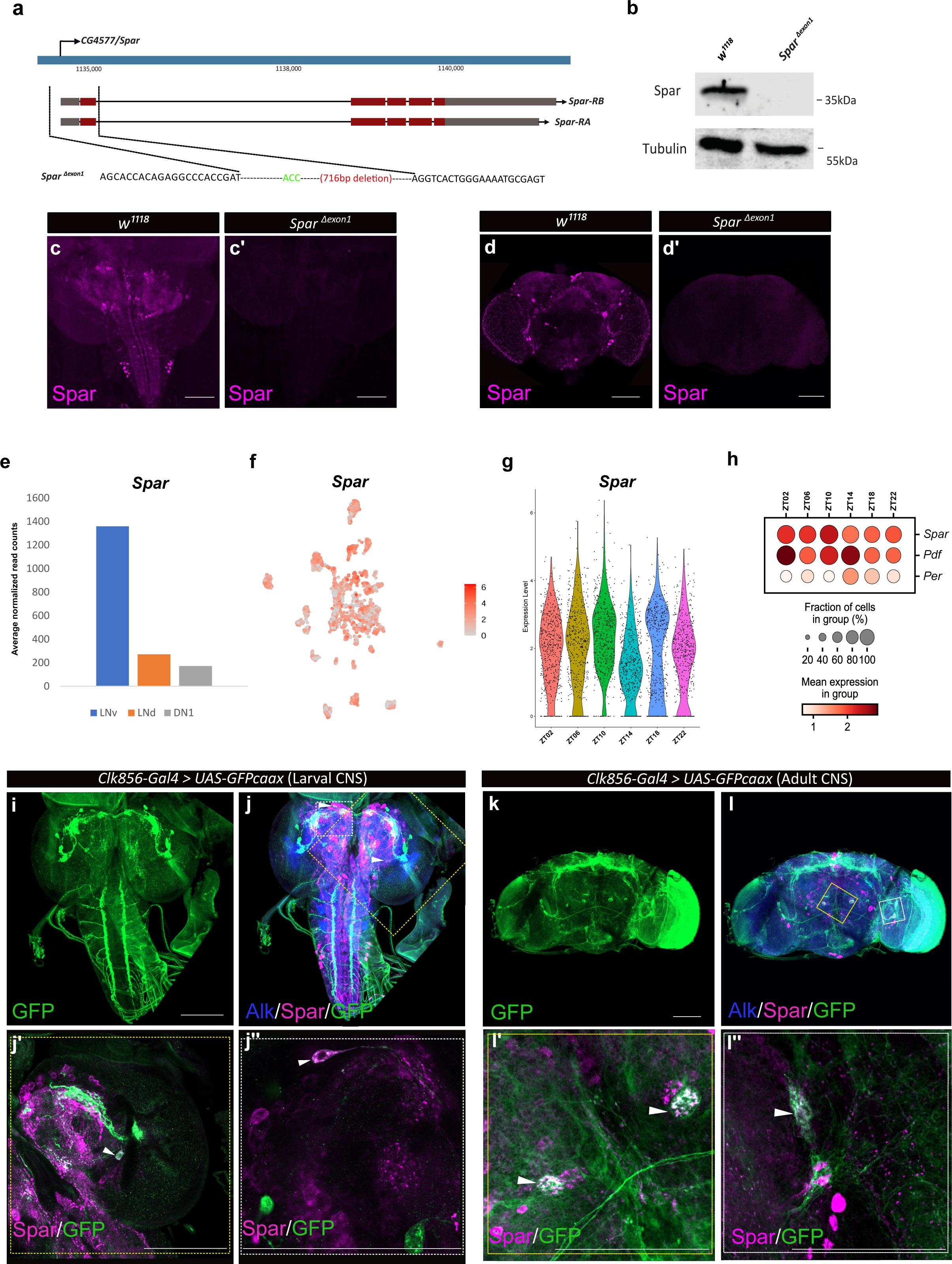
Generation of *Spar^ΔExon1^* mutant and expression of *Spar* in circadian neurons. **a.** Schematic overview of the *Spar* gene locus and the *Spar^ΔExon1^* mutant. Black dotted lines indicate the deleted region, which includes the transcriptional start and exon 1. **b.** Immunoblotting for Spar. Spar protein (35 kDa) is present in larval CNS lysates from wild-type (*w^1118^*) controls but absent in *Spar^ΔExon1^* mutants. **c-d’.** Immunostaining confirms loss of Spar protein expression in the *Spar^ΔExon1^* mutant. Third instar larval **(c-c’)** and adult **(d-d’)** CNS stained for Spar (in magneta). Spar signal is undetectable in *Spar^ΔExon1^*. **e.** Expression of *Spar* in LNv, LNd and DN1 circadian neuronal populations, employing publicly available RNA-seq data (Abruzzi *et al*., 2017). **f.** Feature plot of *Spar* expression in circadian neurons, employing publicly available scRNA-seq data (Ma *et al*., 2021). **g.** Violin plot indicating *Spar* expression throughout the LD cycle, showing light phase (ZT02, ZT06 and ZT10) and dark phase (ZT14, ZT18 and ZT22) expression. **h.** Dotplot comparing *Spar* expression throughout the LD cycle with the previously characterized circadian-associated neuropeptide pigment dispersion factor (*Pdf*) and the core clock gene Period *(per)*. Expression levels and percentage of expressing cells are indicated. **i-j.** Spar expression in clock neurons (*Clk856-Gal4>UAS-GFPcaax*) of the larval CNS **(i-j)**, visualized by immunostaining for Spar (magenta), Alk (in blue) and clock neurons (GFP, in green). **j’-j’’.** Close up of central brain regions (yellow dashed box in **j**) indicating expression of Spar in Clk856-positive neurons (white arrowheads). **k-l.** Immunostaining of *Clk856-Gal4>UAS-GFPcaax* in adult CNS with GFP (in green), Spar (in magenta) and Alk (in blue). **l’-l’’.** Close ups of CNS regions (yellow dashed box in **l**) stained with GFP (in green) and Spar (in red) showing a subset of clock-positive neurons expressing Spar (white arrowheads). Scale bars: 100 μm.

### Spar is expressed in a subset of clock-neurons in the larval and adult CNS

A previous report noted expression of *Spar* in the ventral lateral neuron (LNv), dorsal lateral neuron (LNd) and dorsal neuron 1 (DN1) populations of adult *Drosophila* circadian clock neurons (Abruzzi *et al*, 2017) (**Figure 7e, Table S1)**. A meta-analysis of the publicly available single-cell transcriptomics of circadian clock neurons indicated that almost all adult cluster of clock neurons express *Spar* (Ma *et al*, 2021) **(Figure 7f)**. Additionally, we noted that the expression of *Spar* peaks around Zeitgeber time 10 (ZT10) (coinciding with the evening peak of locomotor activity) **(Figure 7g-h),** although the differences in expression level around the clock with LD or DD cycle were not dramatic **(Figure 7 – figure supplement 2a-c)**. To confirm the expression of Spar in circadian neurons at the protein level we co-stained Spar with a clock neuron reporter (*Clk856-Gal4>UAS-GFP*). A subset of Spar-positive larval CNS neurons appeared to be *Clk856-Gal4>UAS-GFP* positive (**Figure 7i-j’’**). Similarly, a subset of Spar-positive neurons in adults were GFP-positive (**Figure 7k-l’’**), confirming the expression of Spar protein in LNv clock neurons. Taken together, these findings suggest a potential function of the Alk-regulated TaDa-identified target Spar in the maintenance of circadian activity in *Drosophila*.

### *Spar^ΔExon1^* mutants exhibit reduced adult lifespan, activity and circadian disturbances

Given the expression of Spar in circadian neurons of the larval CNS, and the previous observations of a role of Alk mutations in sleep dysregulation in flies (Bai & Sehgal, 2015), we hypothesised that *Spar^ΔExon1^* mutants may exhibit activity/circadian rhythm-related phenotypes. To test this, we first investigated the effects of loss of *Spar* (employing *Spar^ΔExon1^*) and loss of *Alk* (employing a CNS specific loss of function allele of *Alk*, *Alk^ΔRA^* (Pfeifer *et al*., 2022)) on adult lifespan and sleep/activity behaviour using the DAM (*Drosophila* activity monitor) system (Trikinetics Inc.). Both *Alk^ΔRA^* and *Spar^ΔExon1^* mutant flies displayed a significantly reduced lifespan when compared to *w^1118^* controls, with the *Spar^ΔExon1^* group exhibiting a significant reduction in survival at 25 days (**Figure 8a**). Activity analysis in *Alk^ΔRA^* and *Spar^ΔExon1^* flies under 12 h light: 12 h dark (LD) conditions indicated that both *Alk^ΔRA^* and *Spar^ΔExon1^* flies exhibited two major activity peaks, the first centered around Zeitgeber time 0 (ZT0), the beginning of the light phase, the so-called morning peak, and the second around Zeitgeber time 12 (ZT12), the beginning of the dark phase that is called the evening peak (**Figure 8b, black arrows**). Overall activity and sleep profiles per 24 h showed increased activity in *Spar^ΔExon1^* flies (**Figure 8b-d, Figure 8 – figure supplement 1**), that was more prominent during the light phase, with an increase in the anticipatory activity preceding both the night-day and the day-night transition in comparison to *Alk^ΔRA^* and *w^1118^* (**Figure 8b, empty arrows**). Actogram analysis over 30 days showed an increased number of activity peaks in the mutant groups, indicating a hyperactivity phenotype, in comparison to wild-type (**Figure 8d**). Furthermore, mean activity and sleep were also affected; the two mutant groups (*Alk^ΔRA^* and *Spar^ΔExon1^*) displayed significant variations in activity means (**Figure 8e, h-h’; Figure 8 – figure supplement 2**). Analysis of anticipatory activity by quantifying the ratio of activity in the 3 h period preceding light transition relative to activity in the 6 h period preceding light transition as previously described (Harrisingh *et al*, 2007), failed to identify conclusive effects on anticipatory activity in *Spar^ΔExon1^* flies (**Figure 8f-g**). Furthermore, both *Alk^ΔRA^* and *Spar^ΔExon1^* exhibited significant decrease in average sleep during the day per 12 h at young ages (days 5 to 7) (**Figure 8h, Figure 8 – figure supplement 2a)**. In contrast, older flies (days 20 to 22) did not show any significant differences in sleep patterns during the day and per 12 h (**Figure 8h’, Figure 8 – figure supplement 2a’**). The decrease in average sleep in both Alk*^ΔRA^* and *Spar^ΔExon1^* was accompanied by an increase in number of sleep bouts per 12 hours at young age (days 5 to 7) (**Figure 8 – figure supplement 2b**) with no difference in number of sleep bouts at older age (**Figure 8 – figure supplement 2b’**). Rhythmicity analysis showed that *Alk^ΔRA^* and *w^1118^* are more rhythmic in LD compared to *Spar^ΔExon1^* flies **(Figure 8 – figure supplement 3a)**, however when comparing percentage of rhythmic flies among all groups the differences were not significant **(Figure 8 – figure supplement 3a’)**. Moreover, free-running period calculation by Chi-square periodograms showed that both *w^1118^* and *Spar^ΔExon1^* flies exhibit a longer circadian period (higher than 1440 minutes), with 13% of the latter group having a shorter period **(Figure 8 – figure supplement 4a-a’)**. These results demonstrate that Spar is important for normal fly activity and loss of spar affects adult sleep/wake activity.

**Figure 8.**
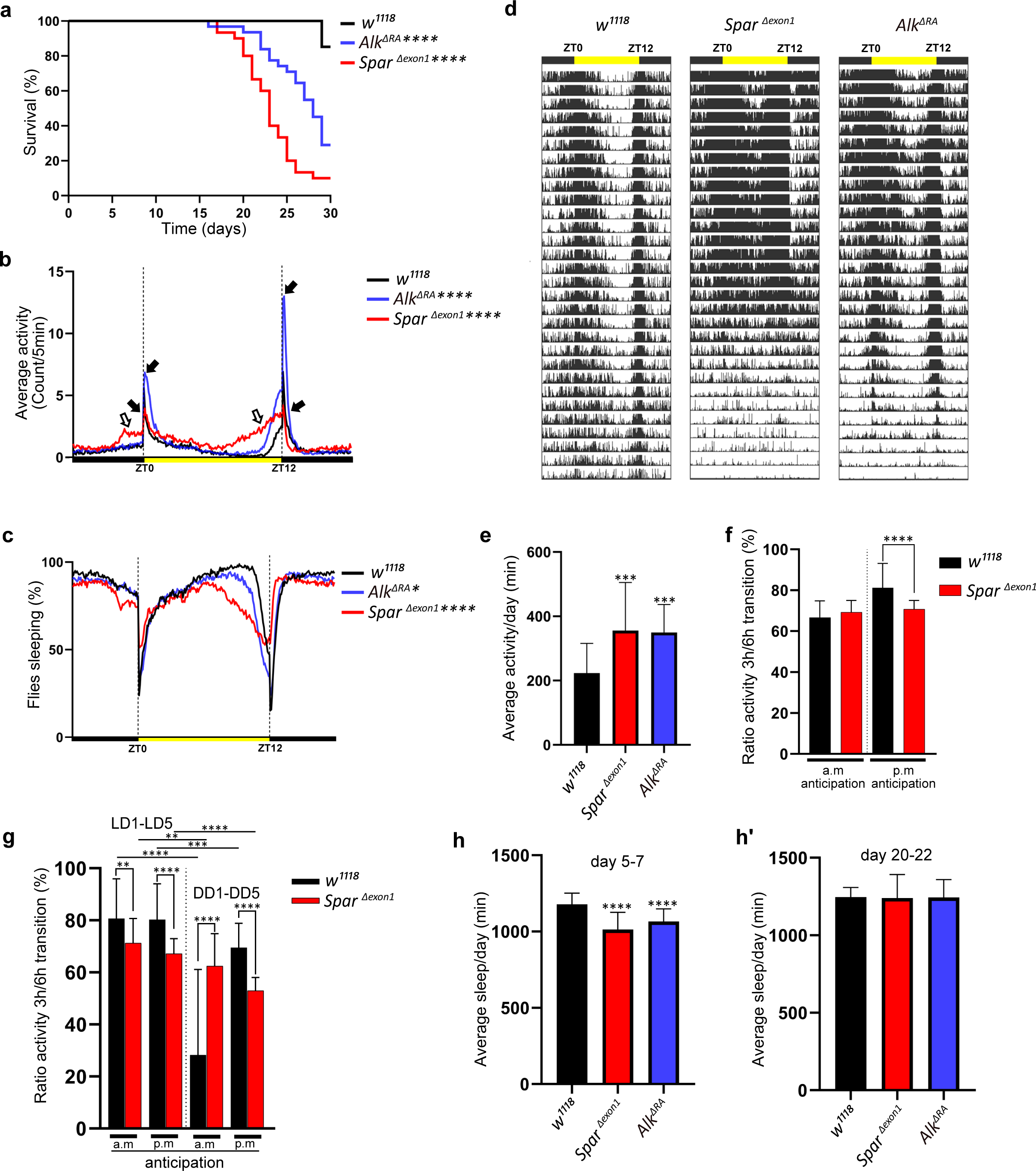
Lifespan and activity plots of *Spar^ΔExon1^* mutants. **a.** Kaplan-Meier survival curve comparing *Alk^ΔRA^* (n=31) and *Spar^ΔExon1^* (n=30) flies to *w^1118^* controls (n=27). Outliers from each group were determined by Tukey’s test, and statistical significance was analyzed by Log-rank Mantel-Cox test (\*\*\*\**p*<0.0001). **b.** Representative activity profile graph illustrating average activity count measured every 5 min across a 24 h span. Black arrows indicate morning and evening activity peaks. Empty arrows indicate anticipatory increase in locomotor activity of *Spar^ΔExon1^* mutant flies occurring before light transition. One-way ANOVA followed by Tukey’s multiple comparisons post-hoc test was used to determine significance between groups (\*\*\*\**p*<0.0001). *w^1118^* (n=27), *Spar^ΔExon1^* (n=30), *Alk^ΔRA^* (n=31). **c.** Representative sleep profile, demonstrating the proportion of flies engaged in sleep measured at 5-minute intervals over a 24-hour period. One-way ANOVA followed by Tukey’s multiple comparisons post-hoc test was used to determine significance between groups (\*\*\*\**p*<0.0001; \**p*<0.05). *w^1118^* (n=27), *Spar^ΔExon1^* (n=30), *Alk^ΔRA^* (n=31). **d.** Representative average actogram of individual flies in each group. Each row corresponds to one day, visualized as 288 bars each representing one 5 min interval. Yellow bar represents the time of the day when the lights are turned on, with ZT0 indicating the morning peak and ZT12 the evening peak. **e.** Mean locomotor activity per day over 30 days. One-way ANOVA followed by Tukey’s multiple comparisons post-hoc test was used to determine significance between groups (\*\*\**p*<0.001). *w^1118^* (n=27), *Spar^ΔExon1^* (n=30), *Alk^ΔRA^* (n=31). **f.** Ratio of the mean activity in the 3 h preceding light transition over the mean activity in the 6h preceding light transition. Activity data is measured over 30 days. a.m. anticipation and p.m. anticipation depict the ratio preceding lights on and lights off respectively. Unpaired student t-test was used to determine the significance between control and *Spar^ΔExon1^* (\*\*\*\**p*<0.0001). *w^1118^* (n=27), *Spar^ΔExon1^* (n=30). **g.** Ratio of the mean activity in the 3 h preceding light transition over the mean activity in the 6h preceding light transition. Activity data is measured over 5 days in LD/DD. a.m. anticipation and p.m. anticipation depict the ratio preceding lights on (or subjective lights on) and lights off (or subjective lights off) respectively. Unpaired student t-test was used to determine the significance between *Spar^ΔExon1^* and controls (*\**\*\*\**p*<0.0001; \*\**p*<0.01); paired student t-test was used to determine significance in each group between the two experimental conditions (\*\*\*\**p*<0.0001; \*\*\**p*<0.001; \*\**p*<0.01). *w^1118^* (n=32), *Spar^ΔExon1^* (n=31). **h-h’.** Mean sleep per day across a 3-day average (Day 5-7 (h), Day 20-22 (h’)). One-way ANOVA followed by Tukey’s multiple comparisons post-hoc test was used to determine significance between groups (\*\*\*\**p*<0.0001). *w^1118^* (n=27), *Spar^ΔExon1^* (n=30), *Alk^ΔRA^* (n=31). The error bars in the bar graphs represents standard deviation.

Since *Spar^ΔExon1^* flies exhibited a hyperactive phenotype during both day and night hours, we sought to investigate a potential role of *Spar* in regulating the endogenous fly clock by assessing fly activity after shift to dark conditions. While control flies adapted to the light-dark shift without any effect on mean activity and sleep, *Spar^ΔExon1^* flies exhibited striking defects in circadian clock regulation (**Figure 9a-b’, Figure 8 – figure supplement 1a-d’**). Comparison of average activity and sleep during 5 days of LD (light-dark) versus 5 days of DD (dark-dark) cycles, identified a reduction in mean activity under DD conditions in *Spar^ΔExon1^* flies (**Figure 9b-b’).** Actogram profiling showed that *Spar^ΔExon1^* flies exhibit a hyperactive profile consistent with our previous data in LD conditions and maintain this hyperactivity when shifted into DD conditions (**Figure 9c, Figure 8 – figure supplement 1**). Further, anticipatory peaks were largely absent on transition to DD cycle in *Spar^ΔExon1^* mutants with no activity peaks observed at either CT0 or at CT12 (**Figure 9b, empty arrows**), consistent with a significant decrease in the a.m. and p.m. anticipatory activity (**Figure 8g)** and altered activity and sleep bouts in these mutants (**Figure 9d, Figure 8 – figure supplement 2**). To confirm that the circadian clock activity defects observed here were specific to loss of *Spar* we conducted a targeted knockdown of *Spar* in clock neurons, employing *Clk856-Gal4. Clk856-Gal4>Spar-RNAi* flies exhibited a significant disruption in both activity and sleep during the DD transition period, consistent with a hyperactivity phenotype (**Figure 9e-g’, Figure 9 – figure supplement 1)**. Further comparison of *Clk856-Gal4>Spar-RNAi* flies relative to controls identified a consistent increase in activity in both LD and DD conditions upon *Spar* knockdown, with a decrease in sleep observed in DD conditions (**Figure 9 – figure supplement 2**). These findings agree with the expression pattern of *Spar* in clock neurons (**Figure 7**), indicating a role for Spar in circadian clock regulation. Rhythmicity analysis comparing LD and DD cycles in *Spar^ΔExon1^* did not show a significant change indicating that *Spar^Δexon1^* flies are mostly rhythmic in LD and DD conditions, whereas as expected, control *w^1118^* flies were less rhythmic in DD conditions (**Figure 9 – figure supplement 3a-a’).** This was also consistent when percentages of rhythmicity were determined, both *w^1118^* and *Spar^Δexon1^* flies were rhythmic **(Figure 9 – figure supplement 3b-b’)**. In terms of circadian period, the majority of *w^1118^* and *Spar^Δexon1^* flies exhibited a longer free running period in DD (**Figure 9 – figure supplement 3c-d)**.

**Figure 9.**
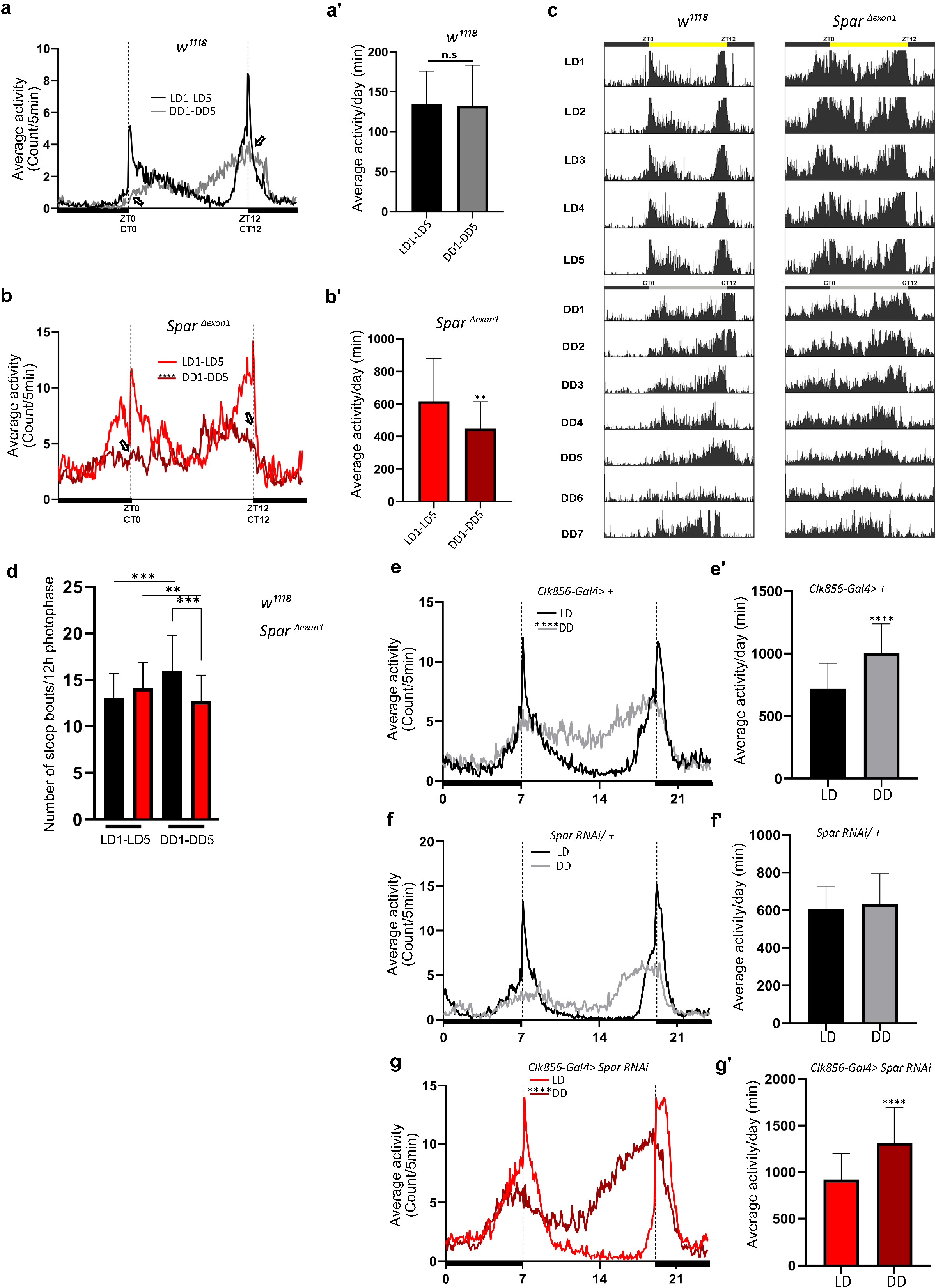
SparΔExon1 mutants exhibit circadian activity disturbances. **a.** Representative activity profile for *w^1118^* controls, illustrating the average activity count measured every 5 min across a 24 h span for Light-Dark (LD) for 5 cycles (black line), subsequently switching to Dark-Dark (DD) for 5 cycles (gray lines). ZT0 and ZT12 represent the start and end of the photoperiod respectively. CT0 and CT12 represent the start and end of the subjective day in constant dark conditions. Empty arrows indicate morning and evening peaks at CT0 and CT12 respectively. Paired student t-test was used to determine significance. *w^1118^* (n=32). **a’**. Mean locomotor activity per day in controls obtained by averaging 5 days in light/dark conditions (LD1-LD5) and 5 days in dark/dark conditions (DD1-DD5). Paired student t-test was used to determine significance. *w^1118^* (n=32) **b**. Representative activity profile graph of *Spar^ΔExon1^* illustrating the average activity count measured every 5 min across 24 h obtained by averaging 5 days in light/dark conditions (LD1-LD5) and 5 days in dark/dark conditions (DD1-DD5). Empty arrows indicate morning and evening peaks at CT0 and CT12 respectively. Paired student t-test was used to determine significance between the two experimental conditions (\*\*\*\**p*<0.0001). *Spar^ΔExon1^* (n=31). **b**’. Mean locomotor activity per day of *Spar^ΔExon1^* obtained by averaging 5 days in light/dark conditions (LD1-LD5) and 5 days in dark/dark conditions (DD1-DD5). Paired student t-test was used to determine significance (\*\*\*\**p*<0.0001). *Spar^ΔExon1^* (n=31). **c**. Representative average actograms of individual *w^1118^* flies (n=32) and *Spar^ΔExon1^* flies (n=31) in LD and DD conditions. Each row corresponds to one day, visualized in 288 bars each representing one 5 min interval. ZT0 and ZT12 represent the start and end of the photoperiod respectively. CT0 and CT12 represent the start and end of the subjective day in constant dark conditions. **d**. Average number of sleep bouts for 12 h photophase over 5 days in LD and the corresponding time over 5 days in DD. Unpaired student t-test was used to determine significance between control (*w^1118^*) and *Spar^ΔExon1^* (\*\*\**p*<0.001). Paired student t-test was used to determine significance between the two experimental conditions (\*\*\**p*<0.001; \*\**p*<0.01). *w^1118^* (n=32), *Spar^ΔExon1^* (n=31). **e**. Representative activity profile graph of *Clk856-Gal4>+* illustrating the average activity count measured every 5 min across a 24 h span for Light-Dark (LD) for 5 cycles (black line) and subsequently switching to Dark-Dark (DD) for 5 cycles (gray lines). Paired student t-test was used to determine significance (\*\*\*\**p*<0.0001). *Clk856-Gal4>+* (n=32). **e**’. Mean locomotor activity per day of *Clk856-Gal4>+* obtained by averaging 5 days in light/dark conditions (LD1-LD5) and 5 days in dark/dark conditions (DD1-DD5). Paired student t-test was used to determine significance (\*\*\*\**p*<0.0001). *Clk856-Gal4>+* (n=32)*. Clk856-GAL4>UAS-Spar RNAi* (n=27). **f**. Representative activity profile graph of *UAS-Spar RNAi*>+ illustrating the average activity count measured every 5 min across 24-hour span obtained by averaging 5 days in light/dark conditions (LD1-LD5) and 5 days in dark/dark conditions (DD1-DD5). A paired student t-test was used to determine the significance between the two experimental conditions. *UAS-Spar RNAi>+* (n=32). **f**’. Graph illustrating the mean locomotor activity per day of *UAS-Spar RNAi*>+ obtained by averaging 5 days in light/dark conditions (LD1-LD5) and 5 days in dark/dark conditions (DD1-DD5). A paired student t-test was used to determine the significance between the two experimental conditions. *UAS-Spar RNAi>+* (n=32). **g**. Representative activity profile graph of *Clk856-Gal4>UAS-Spar RNAi* illustrating the average activity count measured every 5 min across 24 h span obtained by averaging 5 days in light/dark conditions (LD1-LD5) and 5 days in dark/dark conditions (DD1-DD5). Paired student t-test was used to determine significance (\*\*\*\**p*<0.0001). *Clk856-GAL4>UAS-Spar RNAi* (n=27). **g**’. Mean locomotor activity per day for *Clk856-Gal4>UAS-Spar RNAi* obtained by averaging 5 days in light/dark conditions (LD1-LD5) and 5 days in dark/dark conditions (DD1-DD5). Paired student t-test was used to determine the significance (\*\*\*\**p*<0.0001). Error bars represent standard deviation.

## Discussion

With the advent of multiple omics approaches, data integration represents a powerful, yet challenging approach to identify novel components and targets of signaling pathways. The availability of various genetic tools for manipulating Alk signaling in *Drosophila* along with previously gathered omics dataset provides an excellent basis for Alk centered data acquisition. We complemented this with TaDa transcriptional profiling allowing us to generate a rich dataset of Alk-responsive loci with the potential to improve our mechanistic understanding of Alk signaling in the CNS. A striking observation revealed by integrating our TaDa study with scRNAseq data was the enrichment of Alk-responsive genes expressed in neuroendocrine cells. These results are consistent with previous studies reporting expression of Alk in the *Drosophila* larval prothoracic gland (Pan & O’Connor, 2021), the neuroendocrine functions of Alk in mice (Ahmed *et al*., 2022; Reshetnyak *et al*, 2015; Witek *et al*., 2015) and the role of oncogenic ALK in neuroblastoma, a childhood cancer which arises from the neuroendocrine system (Matthay *et al*., 2016; Umapathy *et al*., 2019). In this study, we focused on one target of interest downstream of Alk, however, many additional interesting candidates remain to be explored. These include *CG12594*, *complexin* (*cpx*) and the *vesicular glutamate transporter* (*VGlut*) that also exhibit a high ratio of co-expression with *Alk* in scRNAseq data **(Supplementary** Figure 1). A potential drawback of our TaDa dataset is the identification of false positives, due to non-specific methylation of GATC sites at accessible regions in the genome by Dam protein. Hence, our experimental approach likely more reliably identifies candidates which are downregulated upon Alk inhibition. In our analysis, we have limited this drawback by focusing on genes downregulated upon Alk inhibition and integrating our analysis with additional datasets, followed by experimental validation. This approach is supported by the identification of numerous previously identified Alk targets in our TaDa candidate list.

Employing a strict context dependent filter on our integrated omics datasets identified Spar as a previously uncharacterized Alk regulated neuropeptide precursor. Spar amino acid sequence analysis predicts an N-terminal signal peptide and multiple canonical dibasic PC cleavage sites which are hallmarks of neuropeptide precursors. This is strong indication that Spar is shuttled to the secretory pathway and is post-translationally processed within the Golgi or transport vesicles. Moreover, using mass spectrometry, we were able to identify predicted canonically processed peptides from the Spar precursor in undigested fly brain extracts. While all this points towards a neuropeptide-like function of Spar, other features appear rather unusual for a typical insect neuropeptide. First, the Spar propeptide is quite large for a neuropeptide precursor, and the predicted peptides do not represent paracopies of each other and do not carry a C-terminal amidation signal as is typical for *Drosophila* and other insect peptides (Nässel & Zandawala, 2019; Wegener & Gorbashov, 2008). Moreover, there are no obvious Spar or Spar peptide orthologues in animals outside the Diptera. We noted, however, that Spar is an acidic protein with a pI of 5.1 that lacks any cysteine residue. These features are reminiscent of vertebrate secretogranins, which are packaged and cleaved by PCs and other proteases inside dense vesicles in the regulated secretory pathway in neurosecretory cells (Helle, 2004). Secretogranins have so far not been identified in the *Drosophila* genome (Hart *et al*, 2017). Therefore, the identification of the neurosecretory protein Spar downstream of Alk in the *Drosophila* CNS is particularly interesting in light of previous findings, where VGF (aka secretogranin VII) has been identified as one of the strongest transcriptional targets regulated by ALK in both cell lines and mouse neuroblastoma models (Borenas *et al*., 2021; Cazes *et al*, 2014). *VGF* encodes a precursor polypeptide, which is processed by PCs generating an array of secreted peptide products with multiple functions that are not yet fully understood at this time (Lewis *et al*, 2015; Quinn *et al*, 2021).

Using a newly generated antibody we characterized the expression of Spar in the *Drosophila* CNS, showing that its expression overlaps with the Dimm transcription factor that is expressed in the fly neuroendocrine system (Hewes *et al*., 2003), suggesting that Spar is expressed along with multiple other neuropeptides in pro-secretory cells of the CNS (Park *et al*., 2008). Spar is also expressed in well-established structures such as the mushroom bodies (Crocker *et al*, 2016) (Table 1), which are known to be important in learning and memory and regulate food attraction and sleep (Joiner *et al*, 2006; Pitman *et al*, 2006), and where Alk is also known to function (Bai & Sehgal, 2015; Gouzi *et al*., 2011; Pfeifer *et al*., 2022). Interestingly, Spar is expressed in a subset of peptidergic neurons which emerge from the ventral nerve cord (VNC) and innervate larval body wall muscle number 8. In larvae, these Lk-expressing neurons of the VNC, known as ABLKs, are part of the circuitry that regulates locomotion and nociception, and in adults they regulate water and ion homeostasis (Imambocus *et al*, 2022; Okusawa *et al*, 2014; Zandawala *et al*, 2018). The role of Spar in this context is unknown and requires further investigation. The identity of the Spar receptor, as well as its location, both within the CNS and without, as suggested by the expression of Spar in neurons innervating the larval body wall is another interesting question for a future study. In our current study we focused on characterising Spar in the *Drosophila* CNS. To functionally characterize Spar in this context we generated null alleles with CRISPR/Cas9 and investigated the resulting viable *Spar^ΔExon1^* mutant.

*Spar* transcript expression in *Drosophila* clock neurons has been noted in a previous study investigating neuropeptides in clock neurons, however Spar had not been functionally characterized at the time (Abruzzi *et al*., 2017, Ma *et al*, 2021). We have been able to show that Spar protein is expressed in clock neurons of the larval and adult CNS, findings that prompted us to study the effect of Spar in activity and circadian rhythms of flies. *Drosophila* activity monitoring experiments with *Spar^ΔExon1^* and Alk loss of function (*Alk^ΔRA^*) mutants revealed striking phenotypes in life span, activity and sleep. In *Drosophila* a number of genes and neural circuits involved in the regulation of sleep have been identified (Shafer & Keene, 2021). The role of Alk in sleep has previously been described in the fly, where Alk and the Ras GTPase Neurofibromin1 (Nf1), function together to regulate sleep (Bai and Sehgal, 2015). Indeed, a study in mice has reported an evolutionarily conserved role for Alk and Nf1 in circadian function (Weiss *et al*, 2017). While these studies place Alk and Nf1 together in a signaling pathway that regulates sleep and circadian rhythms, no downstream effectors transcriptionally regulated by the Alk pathway have been identified that could explain its regulation of *Drosophila* sleep/activity. Our data suggest that one way in which Alk signaling regulates sleep is through the control of Spar, as *Spar^ΔExon1^* mutants exhibit a striking activity phenotype. The role of clock neurons and the involvement of circadian input in maintenance of long term memory (LTM) involving neuropeptides such as PDF has been previously described (Inami *et al*, 2022). Since both Alk and Nf1 are also implicated in LTM formation in mushroom body neurons (Gouzi, Bouraimi et al 2018), the potential role of Nf1 in Spar regulation and the effect of Spar loss on LTM will be interesting to test in future work. It can be noted that insulin producing cells (IPCs), DH44 cells of the pars intercerebralis, the Lk producing LHLK neurons of the brain and certain AstA neurons in the brain are involved in regulation of aspects of metabolism and sleep (Barber *et al*, 2021; Cavey *et al*, 2016; Chen *et al*, 2016; Cong *et al*, 2015; Donlea *et al*, 2018; Nässel & Zandawala, 2022; Yurgel *et al*, 2019). Furthermore, the DH44 cells of the pars intercerebralis are major players in regulation feeding and courtship in adults (Barber *et al*., 2021; Cavanaugh *et al*, 2014; Dus *et al*, 2015; King *et al*, 2017; Oh *et al*, 2021).

In conclusion, our TaDa analysis identifies a role for Alk in regulation of endocrine function in *Drosophila*. These results agree with the previously reported broad role of Alk in functions such as sleep, metabolism, and olfaction in the fly and in the hypothalamic-pituitary-gonadal axis and Alk-driven neuroblastoma responses in mice. Finally, we identify *Spar* as the first neuropeptide precursor downstream of Alk to be described that regulates activity and circadian function in the fly.

## Materials and methods

### *Drosophila* stocks and Genetics

Standard *Drosophila* husbandry procedures were followed. Flies were fed on Nutri-Fly® Bloomington Formulation food (Genesee Scientific, Inc.) cooked according to the manufacturer’s instruction. Crosses were reared at 25°C. The following stocks were obtained from Bloomington *Drosophila* Stock Center (BDSC): *w^1118^* (BL3605), *Dimm-Gal4* (also known as *C929-Gal4*) (BL25373), *Clk856-Gal4* (BL93198) and *C155-Gal4* (BL458). The *UAS-Spar RNAi* (v37830) line was obtained from Vienna Drosophila Resource Center. Additional stocks used in this study are the following: *UAS-LT3-NDam-Pol II* (Southall *et al*., 2013), *UAS-Alk^DN^* (*P{UAS-Alk.EC.MYC}* (Bazigou *et al*., 2007)), *UAS-Jeb* (Varshney & Palmer, 2006), *UAS-GFPcaax* (Finley *et al*, 1998), *Alk^Y1335S^* (Pfeifer *et al*., 2022), *Alk^ΔRA^* (Pfeifer *et al*., 2022), *Spar^ΔExon1^* (this study), *UAS-Spar* (this study).

### TaDa Sample preparation

Pan neuronal *C155-Gal4* expressing animals were crossed with either *UAS-LT3-Dam::Pol II* (Control) or *UAS-LT3-Dam::Pol II; UAS-Alk^EC^* (Alk dominant negative sample) and crosses were reared at 25°C. Approximately 100-150 third instar larval brains were dissected in cold PBS for each technical replicate. Genomic DNA was extracted using a QIAGEN blood and tissue DNA extraction kit and methylated DNA was processed and amplified as previously described (Choksi *et al*, 2006; Sun *et al*, 2003) with the following modifications; after genomic DNA extraction, non-sheared gDNA was verified on 1.5% agarose gel, and an overnight DpnI digestion reaction set up in a 50 µl reaction volume. The digestion product was subsequently purified using QIAGEN MiniElute PCR purified Kit and eluted in 50 µl MQ water. 50 µl of DpnI digested and purified DNA was further used for adaptor ligation. Adaptor ligated DNA was amplified using the adaptor specific primer to generate the TaDa-seq library. Amplified DNA from all experimental conditions was repurified (QIAGEN MiniElute PCR purification kit) into 20 µl of MQ water and 200 ng aliquots were run on 1% agarose gel to verify amplification of TaDa library (DNA fragments ranging from 500 bp to 3 kb). The TaDa library was used for PCR-free library preparation followed by paired-end sequencing on an Illumina HiSeq 10x platform (BGI Tech Solutions, Hong Kong).

### TaDa bioinformatics data analysis

TaDa FASTQ paired-end reads of the control sample with three biological replicates and dominant negative samples with two biological replicates (with two technical replicates for both control and dominant negative samples) were obtained for a total of 10 samples and used for subsequent analysis. After base quality assessment, reads were mapped to the Dm6 reference genome of *Drosophila melanogaster* using Bowtie2 (--very-sensitive-local) (Langmead & Salzberg, 2012) and post alignment processes were performed with sam tools and BED tools (Barnett *et al*, 2011; Quinlan, 2014). The *Drosophila melanogaster* reference sequence (FASTA) and gene annotation files were downloaded from Flybase and all GATC coordinates were extracted using fuzznuc (Rice *et al*, 2000) in BED format. Replicates were merged using Sambamba (merge) (Tarasov *et al*, 2015), and fold changes between control and dominant negative samples, obtained by deeptools bamCompare (--centerReads --scaleFactorsMethod readCount --effectiveGenomeSize 142573017 --smoothLength 5 -bs 1) (Ramirez *et al*, 2014) for BIGWIG (BW) file generation. Counts of reads mapped to GATC border fragments were generated using a perl script (GATC_mapper.pl) from DamID-Seq pipeline (Maksimov *et al*, 2016). GATC level counts were converted to gene level counts using Bedtools (intersectBed) (Quinlan, 2014). GATC sites were merged into peaks based on a previous study (Tosti et al., 2018). Log2FC for individual GATC sites were generated using Limma for dominant negative vs control (P < 1e-5) and GATC sites were merged into peaks based on median GATC fragment size in the *Drosophila* genome assembly using mergeWindows (tol=195, max.width=5000) and combineTests function from the csaw package (Lun & Smyth, 2016). Peaks were assigned to overlapping genes and filtered for FDR smaller than 0.05 and mean log2FC less than -2. All peak calling and statistical analysis was performed using the R programming environment. TaDa data can also be visualized using a custom UCSC (University of California, Santa Cruz) Genome Browser session (https://genome-euro.ucsc.edu/s/vimalajeno/dm6). WebGestaltR (Liao *et al*, 2019) was used for GO (Gene Ontology) for significantly downregulated TaDa candidates.

### Integration of TaDa data with scRNA-seq and other omics data

Previously published wild-type third instar larval brain scRNA-Seq data (GSE198850) was employed (Pfeifer *et al*., 2022). Cellular heterogeneity was determined with eight different types of cells, including immature neurons, mature neurons, early neuroblast, NB-enriched cells, NB proliferating cells, optic lobe epithelium (OLE), Repo-positive cells and Wrapper-positive cells. The mature neuron population was divided into two groups for the current study: mature neurons and neuroendocrine cells. The neuroendocrine cell cluster was determined based on canonical markers (Guo *et al*., 2019; Huckesfeld *et al*., 2021; Nässel, 2018; Takeda & Suzuki, 2022; Torii, 2009). Subsequent analysis, including dimensionality reduction/projection or cluster visualization and marker identification was performed using R (Seurat) (Stuart *et al*, 2019) and Python (Scanpy) (Wolf *et al*, 2018) packages. Marker genes for each cluster were identified by FindAllMarkers function (Seurat) (Stuart *et al*., 2019). Clusters were visualized using two-dimensional Uniform Manifold Approximation and Projection (UMAP). The top 500 significantly downregulated genes from TaDa data (FDR<0.05 and mean logFC≤-2) were analysed in the third instar larval brain scRNAseq data. These 500 candidates were used as gene signatures, and signature enrichment analysis carried out using AUCell to determine whether a subset of the input gene set was enriched for each cell (with an enrichment threshold set at >0.196), and the clusters projected in UMAP based on the signature score (AUC score) (Aibar *et al*., 2017). Violin plots, dot plots, feature plots, heatmaps and matrix plots were used to visualize gene expression in the scRNAseq data. Functional enrichment analysis for the common significantly downregulated genes from the TaDa analysis was compared to neuroendocrine cell markers using WebGestaltR (Liao *et al*., 2019).

### Circadian neuron scRNA-Seq data analysis

Publicly available circadian neuron scRNA-Seq data (10x) from the GEO database (GSE157504) was employed to investigate expression of *CG4577* in circadian neurons (Ma *et al*., 2021). The dataset includes two conditions: LD (Light and Dark) and DD (Dark and Dark), as well as six time points: 2 hours, 6 hours, 10 hours, 14 hours, and 22 hours. After preprocessing, 3172 and 4269 cells remained for the LD and DD samples respectively, with a total of 15,743 and 15,461 RNA features. Subsequent analysis, including integration, dimensionality reduction/projection and cluster visualization was performed using R (Seurat) (Stuart *et al*., 2019). Based on clustering, 17 clusters were defined and visualized using two-dimensional Uniform Manifold Approximation and Projection (UMAP). Violin plots, dot plots, and feature plots were employed to visualize gene expression.

### Immunohistochemistry

Relevant tissue (larval CNS or body wall muscle preparation) was dissected in cold PBS and tissues fixed in 4% formaldehyde at 4°C for 1 hour. Samples were washed 3 times with 0.1% PBS Triton X-100, followed by overnight incubation in 4% goat serum, 0.1% PBS Triton X-100. The following primary antibodies were used: guinea pig anti-Alk (1:1000, (Loren *et al*., 2003)), rabbit anti-Alk (1:1000, (Loren *et al*., 2003)), and rabbit anti-Dimm (1:1000 (Allan *et al*, 2005)), chicken anti-GFP (1:1000, Abcam #ab13970), mouse mAb anti-PDF (1:1000, DSHB: C7), rabbit anti-Ilp2 (1:1000, (Veenstra *et al*, 2008)), anti-Dh44 (1:1000, (Cabrero *et al*, 2002)), rabbit anti-AstA (1:3000, (Stay *et al*, 1992) (Vitzthum *et al*, 1996), Jena Bioscience GmbH), rabbit anti-Lk (1:1000, (Cantera & Nässel, 1992)), guinea pig anti-Spar (1:2000, this study), and Alexa Fluor®-conjugated secondary antibodies were from Jackson Immuno Research.

### Image Analysis

Spar fluorescence intensity (Figures 4I and M) was quantified for the minimum complete confocal z-series of each third instar larval brain using Fiji (Schindelin et al, 2012). Confocal images from the 488 nm wavelength channel were analyzed as a Z project. Using a selection tool, Spar positive areas were demarcated, and measurements recorded. Corrected total cell fluorescence (CTCF), in arbitrary units, was measured for each third instar brain as follows: CTCF = integrated density – (area of selected cell × mean fluorescence of background readings) (McCloy RA, et al. Cell Cycle. 2014; Bora P, et al. Commun Biol 2021). Calculated CTCFs were represented in the form of boxplots (n = 12 each for *w^1118^, Alk^Y1255S^, Alk^RA^*. n = 5 each for *C155-Gal4>UAS-GFPcaax* and *C155-Gal4>UAS-Alk^DN^*, n = 7 for *C155-Gal4>UAS-Jeb*).

### Immunoblotting

Third instar larval brains were dissected and lysed in cell lysis buffer (50 mM Tris-Cl, pH7.4, 250 mM NaCl, 1 mM EDTA, 1 mM EGTA, 0.5% Triton X-100, complete protease inhibitor cocktail and PhosSTOP phosphatase inhibitor cocktail) on ice for 20 minutes prior to clarification by centrifugation at 14,000 rpm at 4°C for 15 minutes. Protein samples were then subjected to SDS-PAGE and immunoblotting analysis. Primary antibodies used were: guinea pig anti-Spar (1:1000) (this study) and anti-tubulin (Cell Signaling #2125, 1:20,000). Secondary Antibodies used were: Peroxidase Affinipure Donkey Anti-Guinea Pig IgG (Jackson ImmunoResearch #706-035-148) and goat anti-rabbit IgG (Thermo Fisher Scientific # 32260, 1:5000).

### Generation of anti-Spar antibodies

Polyclonal antibodies against Spar (CG4577) were custom generated in guinea pigs by Eurogentec. Two Spar peptides corresponding to epitopes LQEIDDYVPERRVSS (amino acids 212-226) and PVAERGSGYNGEKYF (amino acids 432-446) of Spar-PA were injected simultaneously.

### Biochemical identification of Spar peptides and phylogenetic analysis

Peptidomic data from our previous study on the role of *Drosophila* carboxypeptidase D (SILVER) in neuropeptide processing (Pauls *et al*., 2019) was re-examined for the occurrence of Spar. Peptides were extracted from brains from 5 d old male flies and analyzed on an Orbitrap Fusion mass spectrometer (Thermo Scientific) equipped with a PicoView ion source (New Objective) and coupled to an EASY-nLC 1000 system (Thermo Scientific). Three (controls) and two (mutants) biological samples (pooled brain extracts from 30 flies) were measured in technical duplicates. The raw data is freely available at Dryad (https://doi.org/10.5061/dryad.82pr5td, for details see (Pauls *et al*., 2019)). Database search was performed against the UniProt Drosophila melanogaster database (UP000000803; 22070 protein entries) with PEAKS XPro 10.6 software (Bioinformatics solution) with the following parameters: peptide mass tolerance: 8 ppm, MS/MS mass tolerance: 0.02 Da, enzyme: “none”; variable modifications: Oxidation (M), Carbamidomethylation (C), Pyro-glu from Q, Amidation (peptide C-term). Results were filtered to 1% PSM-FDR.

To identify Spar precursor sequences in other insects and arthropods, tblastn searches with the PAM30 matrix and a low expectation threshold against the whole *Drosophila* Spar precursor or partial peptides flanked by canonical cleavage sites were performed against the NCBI databank (https://blast.ncbi.nlm.nih.gov/Blast.cgi). The obtained sequences were aligned by the MUSCLE algorithm and plotted using JalView 2(Waterhouse *et al*, 2009).

### CRISPR/Cas9 mediated generation of the *Spar^ΔExon1^* mutant

The *Spar^ΔExon1^* mutant was generated using CRISPR/Cas9 genome editing. Design and evaluation of CRISPR target sites was performed using the flyCRISPR Optimal Target Finder tool (Gratz et al., 2015). Single guide RNA (sgRNA) targeting sequences (sequences available in **Table S1**) were cloned into the pU6-BbsI-chiRNA vector (Addgene, Cat. No. 45946) and injected into *vasa-Cas9* (BDSC, #51323) embryos (BestGene Inc.). Injected flies were crossed to second chromosome balancer flies (BDSC, #9120) and their progeny were PCR-screened for a deletion event. Mutant candidates were confirmed by Sanger sequencing (Eurofins Genomics).

### Generation of *UAS-Spar* fly lines

*UAS-Spar* was generated by cloning (GeneScript) the coding sequence of *CG4577-RA* into EcoRI/XbaI-cut *pUASTattB* vector followed by injection into fly embryos (BestGene Inc.) using attP1 (2^nd^ chromosome, BDSC#8621) and attP2 (3^rd^ chromosome, BDSC#8622) docking sites for phiC31 integrase-mediated transformation. Injected flies were crossed to second or third chromosome balancer flies, and transgenic progeny identified based on the presence of mini-white marker.

### Measurement of pupal size

Late pupae of the indicated genotype were collected and placed on glass slides with double-sided tape. Puparium were imaged with a Zeiss Axio Zoom.V16 stereo zoom microscope with a light-emitting diode ring light and measured using Zen Blue edition software. Both female and male pupae, picked randomly, were used for measurements.

### *Drosophila* activity monitor assay

Up to 32 newly eclosed male flies were transferred into individual glass tubes containing food media (1% agar and 5% sucrose), which were each placed into a DAM2 *Drosophila* activity monitor (Trikinetics Inc). Monitors were then placed in a 25 °C incubator running a 12:12h light:dark cycle, at a constant 60% humidity. Activity was detected by an infrared light beam emitted by the monitor across the center of each glass tube. The experiment was carried out for one month, and the raw binary data was acquired by the DAMSystem310 software (Trikinetics Inc.). The LD/DD experiment was performed according to previously published work (Chiu *et al*, 2010); adult flies were first entrained for 5 days in normal light:dark cycle and on the last day (LD5), the light parameters were switched off and flies were then conditioned in complete dark:dark settings for 7 days. Raw data analysis was carried out using a Microsoft Excel macro (Berlandi *et al*, 2017) taking into consideration 5 min of inactivity as sleep and more than 24 h of immobility as a death event. The activity and sleep parameters are calculated for each day of the experiment as average from data of all living animals at this time and are displayed over the duration of the experiment. Calculation of anticipatory activity was performed accordingly to previously published work (Harrisingh *et al*., 2007) by quantifying the ratio of activity in the 3h preceding light transition to activity in the 6h preceding light transition, defined as a.m and p.m aniticipation for the 6h period before lights-on and 6h period before lights-off, respectively. Actogram activity profile charts were generated using ActogramJ 1.0 (https://bene51.github.io/ActogramJ/index.html) and ImageJ software (https://imagej.nih.gov/ij/). ActogramJ was further used to generate the chi-square periodogram for each single fly in order to calculate the power value of rhythmicity and the percentage of rhythmic flies. All statistical analysis were performed using GraphPad Prism 8.4.2.

## Data visualization and schematics

Schematics were generated at Biorender.com and Bioicons.com. The pipeline icon by Simon Dürr https://twitter.com/simonduerr is licensed under CC0 https://creativecommons.org/publicdomain/zero/1.0/. Boxplots in Figure 3d and Figure 7 – figure supplement 1 were generated using BoxplotR (http://shiny.chemgrid.org/boxplotr/). Boxplots in Figure 4 were generated using GraphPad Prism 9.

## Datasets used in this study

| Dataset name | Brief description | Reference | NCBI gene Expression omnibus ID/database IDs |
| --- | --- | --- | --- |
| Targeted DamID (TaDa) dataset | Pol II occupancy of control (*UAS-Dam-Pol II*) and *Alk* knockdown (*UAS-Dam-Pol II; Alk<sup>DN</sup>*) conditions in 3rd instar *Drosophila* larval CNS. | This study | GSE229518 |
| Bulk RNA-seq dataset - *Drosophila Alk* alleles | Transcriptomes of *Alk* gain-of-function (*Alk<sup>Y1355S</sup>*), *Alk* CNS specific loss-of-function (*Alk<sup>RA</sup>*) and control (*w<sup>1118</sup>*) 3rd instar *Drosophila* larval CNS. | Pfeifer *et al.*, 2022 | GSE198812 |
| 3rd instar Larval CNS Single cell RNA-seq dataset | Single cell transcriptome of control (*w<sup>1118</sup>*) *Drosophila* 3rd instar larval CNS. | Pfeifer *et al.*, 2022 | GSE198850 |
| 1st instar larval CNS Single cell RNA-seq dataset | Single cell transcriptome of control (*Canton-S*) *Drosophila* 1st instar larval CNS. | Brunet Avalos *et al.*, 2019 | GSE134722 |
| Adult CNS single cell RNA-seq dataset | Single cell transcriptome of control (*w<sup>1118</sup>*) *Drosophila* adult brains. | Davie *et al.*, 2018 | GSE107451 |
| Adult CNS peptidomics data | Peptidomic mass spectrometry analysis of control (*FM7h;hs-svr*) and SILVER processing mutant (*svr<sup>PG33</sup>;hs-svr*) in *Drosophila* adult brains. | Pauls *et al.*, 2019 | Dryad:<br>https://doi.org/10.5061/dryad.82pr5td |
| Bulk RNA-seq dataset - *Drosophila* circadian clock neurons | Transcriptome of circadian clock neurons from *Drosophila* adult brains. | Abruzzi *et al.*, 2017 | GSE77451 |
| Single cell RNA-seq dataset - *Drosophila* circadian clock neurons | Single cell transcriptome of circadian clock neurons (*Clk856-Gal4> EGFP*) in *Drosophila* adult brains. | Ma *et al.*, 2021 | GSE157504 |

## Acknowledgements

The authors thank Jonathan Benito Sipos and Stefan Thor for the kind gift of anti-Dimmed antibodies, as well as Jan Veenstra for kindly gifting anti-Dh44 and anti-Ilp2. C7 anti-PDF (developed by J. Blau) was obtained from the Developmental Studies Hybridoma Bank, created by the NICHD of the NIH and maintained at The University of Iowa, Department of Biology, Iowa City, IA 52242. We acknowledge Bloomington Drosophila Stock Center (NIH P40OD018537) for fly stocks used in this study. We thank Hisae Mori for providing support for fly lab maintenance. We thank members of the Palmer, Hallberg lab and Anne Uv for critical feedback on the manuscript. This work has been supported by grants from the Swedish Cancer Society (RHP CAN21/01549), the Children’s Cancer Foundation (RHP 2019-0078), the Swedish Research Council (RHP 2019-03914), the Swedish Foundation for Strategic Research (RB13-0204), the Göran Gustafsson Foundation (RHP2016) and the Knut and Alice Wallenberg Foundation (KAW 2015.0144). MS and JS are supported by the Medical Practice Plan (MPP) at the American University of Beirut.

Author contributions: SKS and RHP conceived the research. Wet lab experiments were conducted by SKS, LM, JS, MK, GU, PM-G, and TM. VA carried out all bioinformatics analysis. LM assisted with *Spar* mutant generation, validation, and immunohistochemistry. JS and MK performed and analyzed all *Drosophila* activity monitoring experiments under the supervision of MS. GU performed immunoblotting. PM-G assisted in optimizing and performing TaDa experiments, and TM performed image analysis and assisted with immunohistochemistry. AS and CW analyzed mass spectrometry peptidomic data and performed the phylogenetic analysis. DN provided critical feedback and design input. RHP, CW and MS supervised the project. SKS and RHP wrote the first manuscript draft that was further developed with all authors.

## Competing interests

The authors declare that they have no competing interests

## Data Availability Statement

The original contributions presented in the study are included in the article/Supplementary Material. The TaDa dataset has been deposited in Gene Expression Omnibus (GEO) under the accession number GSE229518. The genome browser tracks for the TaDa peak analysis can be found at: https://genome-euro.ucsc.edu/s/vimalajeno/dm6.

## Supplementary figure legends

**Figure 1 – figure supplement 1. TaDa third instar larval CNS sample validation and additional data analysis. a-b’.** Expression of mCherry in the larval CNS reflects Dam-PolII expression. Third instar larval brains were stained for Alk (in green) and mCherry (in red) confirming expression of Dam-PolII in the TaDa system. Scale bars: 100 μm. **c.** Schematic overview of the TaDa analysis experimental workflow. Brains from third instar wandering larvae were dissected, and methylated DNA digested with Dpn1 restriction endonuclease. The resulting DNA fragment library was amplified, sequenced and analysed through TaDa bioinformatics pipelines. **d.** Bar graph showing total number of reads in each replicate of the TaDa dataset. **d’.** Bar graph showing percentage of reads aligned to *Drosophila* genome in each replicate. **e.** Correlation plot of samples (*control* and *Alk^DN^*) and replicates shows no significant intra-replicate differences. **f.** Line graph indicating the relative distance to TSS of different samples compared to random regions. **g.** Pol ll occupancy profile of *Alk* in *Alk^DN^* compared to control indicates a higher pol II occupancy in exons 1 to exon 7, in agreement with the expression of the *Alk^DN^* transgene.

**Figure 2 – figure supplement 1. a-a’’.** Alk staining in *Dimm-Gal4>UAS-GFPcaax* third instar larval CNS confirms Alk expression in Dimm-positive cells. Alk (in magenta) and GFP (in green), close-ups indicating overlapping cells (indicated by yellow arrowheads) in the larval ring gland corpora cardiaca cells **(b-b’’)** central brain **(c-c’’)** and ventral nerve cord **(d-d’’)**. Scale bars: 100 μm.

**Figure 3 – figure supplement 1. Co-expression of *Alk* and *Spar* in publicly available *Drosophila* CNS scRNA-seq datasets.** UMAP showing co-expression of *Alk* and *CG4577* in different cell clusters in publicly available scRNA-seq datasets (Brunet Avalos *et al*., 2019) from first instar larval CNS **(a)** and **(b)** adult CNS (Davie *et al*., 2018). **c.** Pairwise alignment of CG4577-PA and CG4577-PB showing isoform-specific differences at amino acid positions 405 and 406 (highlighted in yellow).

**Figure 3 – figure supplement 2. Alignment of *CG4577* orthologs in flies (Brachycera). a.** Alignment of *Drosophila melanogaster CG4577* orthologs in the family *Drosophilidae* (vinegar flies, including the fruit fly *Drosophila melanogaster)*. **b.** Alignment of *Drosophila melanogaster CG4577* orthologs in other brachyceran taxa.

**Figure 4 – figure supplement 1. a-a’’** Immunostaining of *w^1118^* third instar larval brains with Spar (green) and Alk (magenta) revealing overlapping expression (indicated by yellow arrowheads) in central brain, ring gland corpora cardiaca and ventral nerve cord. Close-ups indicating overlapping cells in central brain **(b-b’’**, corresponding to dashed box on left in A) and ring gland corpora cardiaca cells **(c-c’’**, corresponding to dashed box on right in A). Scale bars: 100 μm.

**Figure 4 – movie supplement 1.** Z-stack projection video of Figure 4e’

**Figure 4 – movie supplement 2.** Z-stack projection video of Figure 4e’’

**Figure 4 – figure supplement 2. a-a’’** Adult CNS showing Spar expression in Dimm-positive cells. Spar (in magenta) and Dimm (in green), close-ups **(b-b’’)** indicated by boxed region and white arrows indicating representative co-expressed markers cells. Scale bars: 100 μm.

**Figure 6 – figure supplement 1. *Spar* expression in adult neuropeptide expressing neuronal populations**. **a.** Immunostaining of *w^1118^* adult CNS with anti-Spar (in magenta) and anti-PDF (in green). Closeups (**b-b’’)** of PDF- and Spar-positive LNv neurons, indicated by white arrowheads. **c.** Immunostaining of *w^1118^* adult CNS with Spar (in magenta) and Dh44 (in green). Closeups (**d-d’’)** of Dh44- and Spar-positive neurons, indicated by white arrowheads. **e.** Immunostaining of *w^1118^* adult CNS with Spar (in magenta) and Ilp2 (in green). Closeups (**f-f’’)** showing the close proximity and overlap of Ilp2-positive and Spar-positive neurons in central brain, indicated by white arrowheads. **g.** Immunostaining of *w^1118^* adult CNS with Spar (in magenta) and AstA (in green). Closeups (**h-h’’)** showing AstA- and Spar-positive neurons in central brain indicated by white arrowheads. Scale bars: 100μm.

**Figure 7 – figure supplement 1. Spar does not affect the Alk-regulated pupal size phenotype.** Overexpression of Spar (*C155-Gal4>UAS-Spar*) or *Spar RNAi* (*C155-Gal4>UAS-Spar RNAi*) in CNS does not significantly affect pupal size compared to previously characterized controls such as *Alk^DN^* (*C155-Gal4>UAS-Alk^DN^*), which significantly increases pupal size and overexpression of *Jeb* (*C155-Gal4>UAS-Alk^DN^*), which significantly decreases pupal size compared to controls (*C155-Gal4>+*) (n.s = not significant, \*\**p*<0.05, \*\*\**p*<0.01). Center lines in boxplots indicate medians; box limits indicate the 25th and 75th percentiles; whiskers extend 1.5 times the interquartile range from the 25th and 75th percentiles, crosses represent sample means; grey bars indicate 83% confidence intervals of the means; data points are plotted as grey circles.

**Figure 7 – figure supplement 2. *Spar* expression in circadian neuronal clusters. a-b.** Feature plots depicting the expression of *Spar* in publicly available circadian neuronal scRNA-seq data (Ma *et al*., 2021) throughout the LD cycle (Zeitgeber time) (**a**) and DD cycle (Circadian time) **(b). c.** Dotplot showing *Spar* expression throughout the DD cycle along with the previously characterized circadian associated neuropeptide *Pdf* and the core clock gene *Per.* Peak expression of *Spar* and *Per* is observed at CT10.

**Figure 8 – figure supplement 1. a.** Representative activity profile graph of control (*w^1118^*) and *Spar^ΔExon1^* illustrating average activity count measured every 5 min across a 24 h span obtained by averaging 5 days in light/dark conditions (LD1-LD5). Unpaired student t-test was used to determine significance (\*\*\*\**p*<0.0001). *w^1118^* (n=32), *Spar^ΔExon1^* (n=31)**. a’.** Mean locomotor activity per day of control and *Spar^ΔExon1^* obtained by averaging 5 days in light/dark conditions (LD1-LD5). Unpaired student t-test was used to determine significance (\*\*\*\**p*<0.0001). **b.** Representative activity profile graph of control and *Spar^ΔExon1^* illustrating the average activity count measured every 5 min across a 24 h span obtained by averaging 5 days in dark/dark conditions (DD1-DD5). CT0 and CT12 represent the start and end of the subjective day in constant dark conditions respectively. Unpaired student t-test was used to determine significance (\*\*\*\**p*<0.0001). *w^1118^* (n=32), *Spar^ΔExon1^* (n=31)**. b’.** Mean locomotor activity per day for control and *Spar^ΔExon1^* obtained by averaging 5 days in dark/dark conditions (DD1-DD5). Unpaired student t-test was used to determine significance (\*\*\*\**p*<0.0001). *w^1118^* (n=32), *Spar^ΔExon1^* (n=31)**. c.** Representative sleep profile graph of control and *Spar^ΔExon1^* illustrating the average activity count measured every 5 min across a 24 h span obtained by averaging 5 days in light/dark conditions (LD1-LD5). Unpaired student t-test was used to determine significance (\*\*\*\**p*<0.0001). **c’.** Graph illustrating mean sleep per day of control and *Spar^ΔExon1^* obtained by averaging 5 days in light/dark conditions (LD1-LD5). Unpaired student t-test was used to determine significance (\*\*\*\**p*<0.0001). *w^1118^* (n=32), *Spar^ΔExon1^* (n=31)**. d.** Representative sleep profiles of controls and *Spar^ΔExon1^* illustrating the average activity count measured every 5 min across a 24 h span obtained by averaging 5 days in dark/dark conditions (DD1-DD5). Unpaired student t-test was used to determine significance (\*\*\*\**p*<0.0001). *w^1118^* (n=32), *Spar^ΔExon1^* (n=31)**. d’.** Mean sleep per day of control and *Spar^ΔExon1^* obtained by averaging 5 days in dark/dark conditions (DD1-DD5). Unpaired student t-test was used to determine significance (\*\*\*\**p*<0.0001). *w^1118^* (n=32), *Spar^ΔExon1^* (n=31). Error bars represent standard deviation.

**Figure 8 – figure supplement 2. a-a’.** Mean sleep per 12 h photophase over 3 days (Day 5-7 (**a**), Day 20-22 (**a’**)). One-way ANOVA followed by Tukey’s multiple comparisons post-hoc test was used to determine significance groups (\*\*\*\**p*<0.0001). *w^1118^* (n=27), *Spar^ΔExon1^* (n=30), *Alk^ΔRA^* (n=31). **b-b’.** Graph illustrating the average number of sleep bouts for 12 h photophase over 3 days (Day 5-7 (**b**), Day 20-22 (**b’**)). A one-way ANOVA followed by Tukey’s multiple comparisons post-hoc test was used to determine the significance between groups (\*\*\*\**p*<0.0001)*. w^1118^* (n=27), *Spar^ΔExon1^* (n=30), *Alk^ΔRA^* (n=31). Error bars represent standard deviation.

**Figure 8 – figure supplement 3. *Spar^ΔExon1^* flies retain a hyperactive profile when shifted to dark/dark conditions. a.** Graph illustrating the Qp statistical value (rhythmicity power) obtained by generating Chi-square periodograms of control (*w^1118^*), *Spar^ΔExon1^*, and *Alk^ΔRA^* flies. One-way ANOVA followed by Tukey’s multiple comparisons post-hoc test was used to determine significance between groups (\*\*\*\**p*<0.0001). *w^1118^* (n=27), *Spar^ΔExon1^* (n=30), *Alk^ΔRA^* (n=31). **a’.** Representative graph of percentage of rhythmicity of *w^1118^*, *Spar^ΔExon1^*, and *Alk^ΔRA^* flies. *w^1118^* (n=27), *Spar^ΔExon1^* (n=30), *Alk^ΔRA^* (n=31). Error bars represent standard deviation.

**Figure 8 – figure supplement 4. a.** Graph illustrating the percentage of arrhythmic flies, and flies with a period higher or lower than 1440 minutes (24 h). Flies were maintained under LD conditions and 14 days were selected to calculate the period by generating Chi-Square periodograms for each fly in the group. Only data from rhythmic flies was selected to calculate the percentage of flies having a higher or lower period than 1440 minutes. **a’**. Average periods of flies over 14 days in LD conditions. Flies with an arrhythmic profile were not selected for statistical analysis. Unpaired student t-test was used to determine significance between the two groups. Error bars represent standard deviation.

**Figure 9 – figure supplement 1. a.** Representative sleep profile graph of *Clk856-Gal4>+* illustrating the percentage of time sleeping measured every 5 min across a 24 h span obtained by averaging 5 days in light/dark conditions (LD1-LD5) and 5 days in dark/dark conditions (DD1-DD5). Paired student t-test was used to determine significance (\*\*\*\**p*<0.0001). *Clk856-Gal4>+ (n=32).* **a**’. Mean sleep per day of *Clk856-Gal4>+* obtained by averaging 5 days in light/dark conditions (LD1-LD5) and 5 days in dark/dark conditions (DD1-DD5). Paired student t-test was used to determine significance (\*\*\*\**p*<0.0001). *Clk856-Gal4>+ (n=32)*. **b**. Representative sleep profile of *UAS-Spar RNAi*>+ illustrating the percentage of time that flies spend sleeping measured every 5 min across a 24-hour span obtained by averaging 5 days in light/dark conditions (LD1-LD5) and 5 days in dark/dark conditions (DD1-DD5). Paired student t-test was used to determine the significance between the two experimental conditions (*\*\*\*\*p*<0.0001). *UAS-Spar RNAi/*+ (n=32). b’. Mean sleep per day of *UAS-Spar RNAi/*+ obtained by averaging 5 days in light/dark conditions (LD1-LD5) and 5 days in dark/dark conditions (DD1-DD5). Paired student t-test was used to determine the significance between the two experimental conditions (*\*\*\*\*p*<0.0001). *UAS-Spar RNAi/*+ (n=32). c. Representative sleep profile graph of *Clk856-Gal4>Spar RNAi* illustrating the percentage of time sleeping measured every 5 min across a 24 h span obtained by averaging 5 days in light/dark conditions (LD1-LD5) and 5 days in dark/dark conditions (DD1-DD5). Paired student t-test was used to determine significance (\*\*\*\**p*<0.0001). *Clk856-Gal4>UAS-Spar RNAi* (n=27). c’. Mean sleep per day of *Clk856-Gal4>Spar RNAi* obtained by averaging 5 days in light/dark conditions (LD1-LD5) and 5 days in dark/dark conditions (DD1-DD5). Paired student t-test was used to determine significance (\*\*\*\**p*<0.0001). *Clk856-Gal4>UAS-Spar RNAi* (n=27). Error bars represent standard deviation.

**Figure 9 – figure supplement 2. a.** Representative activity profile graph of *UAS-Spar RNAi* driven by *Clk856-Gal4* and the two control groups *Clk856-Gal4*>+ and *UAS-Spar RNAi/*+ illustrating the average activity count measured every 5 min across 24-hour obtained by averaging 5 days in light/dark conditions (LD1-LD5). One-way ANOVA followed by Tukey’s multiple comparisons post-hoc test was used to determine the significance between groups (*\*p*<0.05, *\*\*\*p*<0.001). *Clk856-Gal4>+* (n=32), *Clk856-Gal4>UAS-Spar RNAi* (n=27), and *UAS-Spar RNAi/+* (n=32). **a’**. Graph illustrating mean locomotor activity per day of *Clk856-Gal4*>+, *Clk856-Gal4*>*UAS-Spar RNAi* and *UAS-Spar RNAi/+* obtained by averaging 5 days in light/dark conditions (LD1-LD5). One-way ANOVA followed by Tukey’s multiple comparison post-hoc test was used to determine the significance between groups (*\*\*\*p*<0.001, *\*\*\*\*p*<0.0001). *Clk856-Gal4>+* (n=32), *Clk856-GAL4>UAS-Spar RNAi* (n=27), and *UAS-Spar RNAi/+* (n=32). **b**. Representative activity profile graph of *UAS-Spar RNAi* driven by *Clk856-Gal4* and the two control groups *Clk856-Gal4*>+ and *UAS-Spar RNAi/*+ illustrating the average activity count measured every 5 min across a 24-hour span obtained by averaging 5 days in dark/dark conditions (DD1-DD5). One-way ANOVA followed by Tukey’s multiple comparison post-hoc test was used to determine the significance between groups (*\*\*\*\*p*<0.0001). *Clk856-Gal4>+* (n=32), *Clk856-Gal4>UAS-Spar RNAi* (n=27), and *UAS-Spar RNAi/+* (n=32). **b’**. Graph illustrating the mean locomotor activity per day of *Clk856-Gal4*>+, *Clk856-Gal4*>*UAS-Spar RNAi* and *UAS-Spar RNAi/+* obtained by averaging 5 days in dark/dark conditions (DD1-DD5). One-way ANOVA followed by Tukey’s multiple comparison post-hoc test was used to determine the significance between groups (*\*\*\*\*p*<0.0001). *Clk856-Gal4>+* (n=32), *Clk856-Gal4>UAS-Spar RNAi* (n=27), and *UAS-Spar RNAi/+* (n=32). **c**. Representative sleep profile graph of *UAS-Spar RNAi* driven by *Clk856-Gal4* and the two control groups *Clk856-Gal4*>+ and *UAS-Spar RNAi*/+ illustrating the average activity count measured every 5 min across a 24-hour span obtained by averaging 5 days in light/dark conditions (LD1-LD5). One-way ANOVA followed by Tukey’s multiple comparison post-hoc test was used to determine the significance between groups (*\*\*\*p*<0.01). *Clk856-Gal4>+* (n=32), *Clk856-GAL4>UAS-Spar RNAi* (n=27), and *UAS-Spar RNAi>+* (n=32). **c’**. Graph illustrating the mean sleep per day of *Clk856-Gal4*>+, *Clk856-Gal4*>*UAS-Spar RNAi* and *UAS-Spar RNAi/+* obtained by averaging 5 days in light/dark conditions (LD1-LD5). One-way ANOVA followed by Tukey’s multiple comparison post-hoc test was used to determine the significance between groups (*\*\*\*\*p*<0.0001). **d**. Representative sleep profile graph of *UAS-Spar RNAi* driven by *Clk856-Gal4* and the two control groups *Clk856-Gal4*>+ and *UAS-Spar RNAi*/+ illustrating the average activity count measured every 5 min across a 24-hour span obtained by averaging 5 days in dark/dark conditions (DD1-DD5). One-way ANOVA followed by Tukey’s multiple comparison post-hoc test was used to determine the significance between groups (* *p*<0.005, *\*\*\*\*p*<0.0001). *Clk856-Gal4>+* (n=32), *Clk856-Gal4>UAS-Spar RNAi* (n=27), and *UAS-Spar RNAi/+* (n=32). **d’**. Graph illustrating the mean sleep per day of *Clk856-Gal4*>+, *Clk856-Gal4*>*UAS-Spar RNAi* and *UAS-Spar RNAi*/+ obtained by averaging 5 days in dark/dark conditions (DD1-DD5). One-way ANOVA followed by Tukey’s multiple comparisons post-hoc test was used to determine the significance between groups (*\*p*<0.05, *\*\*\*\*p*<0.0001). *Clk856-Gal4>+* (n=32), *Clk856-Gal4>UAS-Spar RNAi* (n=27), and *UAS-Spar RNAi/+* (n=32). Error bars represent standard deviation.

**Figure 9 – figure supplement 3. a.** Qp statistical value obtained by generating Chi-square periodograms of control (*w^1118^*) flies in 5 days LD and 7 days DD conditions. Paired student t-test was used to determine significance (\*\*\**p*<0.001). *w^1118^* (n=32). **a**’. Qp statistical value obtained by generating Chi-square periodograms of *Spar^ΔExon1^* flies in 5 days LD and 7 days DD conditions. Paired student t-test was used to determine significance. *Spar^ΔExon1^* (n=31). **b.** Percentage rhythmicity of control (*w^1118^*) flies in LD vs DD conditions. *w^1118^* (n=32). **b**’. Percentage rhythmicity of *Spar^ΔExon1^* flies in LD vs DD conditions. *Spar^ΔExon1^* (n=31). **c.** Percentage of flies with a period higher or lower than 1440 minutes (24 h). Flies were maintained for 5 days under LD conditions and shifted to 7 days under DD (free-run). The period was calculated by generating Chi-Square periodograms for each fly in the group. Only data from rhythmic flies was selected to calculate the percentage of flies having a period higher or lower than 1440 minutes. *w^1118^* (n=32), *Spar^ΔExon1^* (n=31). d. Graph illustrating the average periods of flies over 5 days in LD and 7 days in DD (free-run). Flies with an arrhythmic profile were not selected for the statistical analysis. An unpaired student t-test was used to determine the significance between the two groups (\*\**p*<0.01). *w^1118^* (n=32), *Spar^ΔExon1^* (n=31). Error bars represent standard deviation.

## Additional Supplementary information

**Supplementary Figure 1.** Feature plots visualizing expression of TaDa-identified genes expressed in neuroendocrine cells in scRNA-seq from third instar larval CNS (Pfeifer *et al*., 2022). TaDa candidates *CG12594, cpx* and *VGlut* are shown.

**Supplementary Table 1.** TaDa data Alk^DN^ downregulated genes (Sheet 1). RNASeq normalized read count data of *CG4577* in control (*w^1118^*), *Alk^RA^* and *Alk^Y1355S^* conditions (Sheet 2). RNAseq average normalized read count data of *Spar* in LNv, LNd and DN1 clock neuronal cells (Sheet 3). *Spar^ΔExon1^* mutant CRISPR single guide RNA target and screening primer information (Sheet 4).

## Notes

### Competing Interest Statement

The authors have declared no competing interest.

### Summary of Updates

Minor corrections have been included in this version of the manuscript.These corrections can be found in Figs 8 and 9.

