## Supplementary material for "The Alk receptor tyrosine kinase regulates Sparkly, a novel activity regulating neuropeptide precursor in the *Drosophila* CNS": Supp Figures

*Sukumar et al., 2024* - **Figure 1 - figure supplement 1**

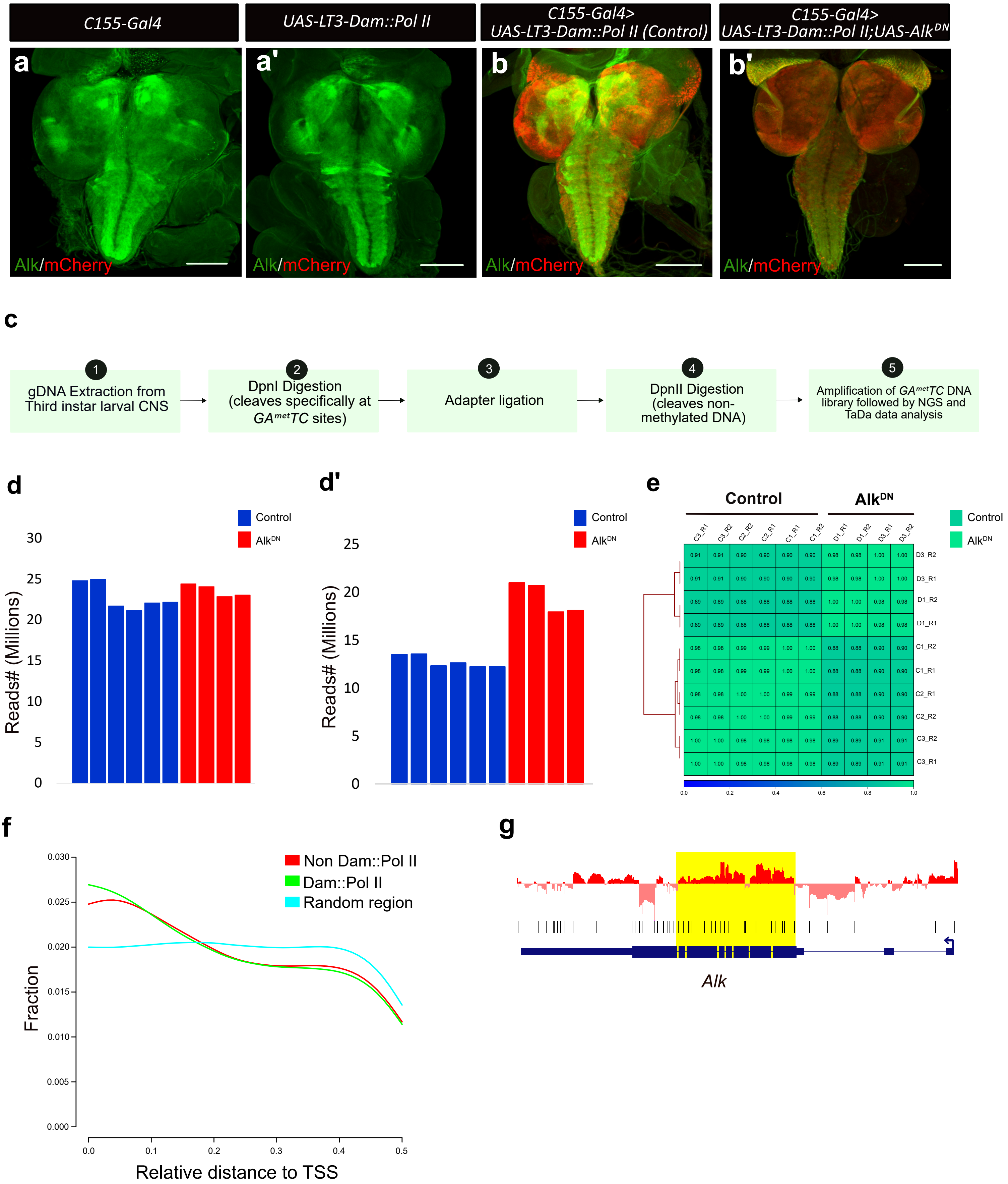

*Sukumar et al., 2024* - **Figure 2 - figure supplement 1**

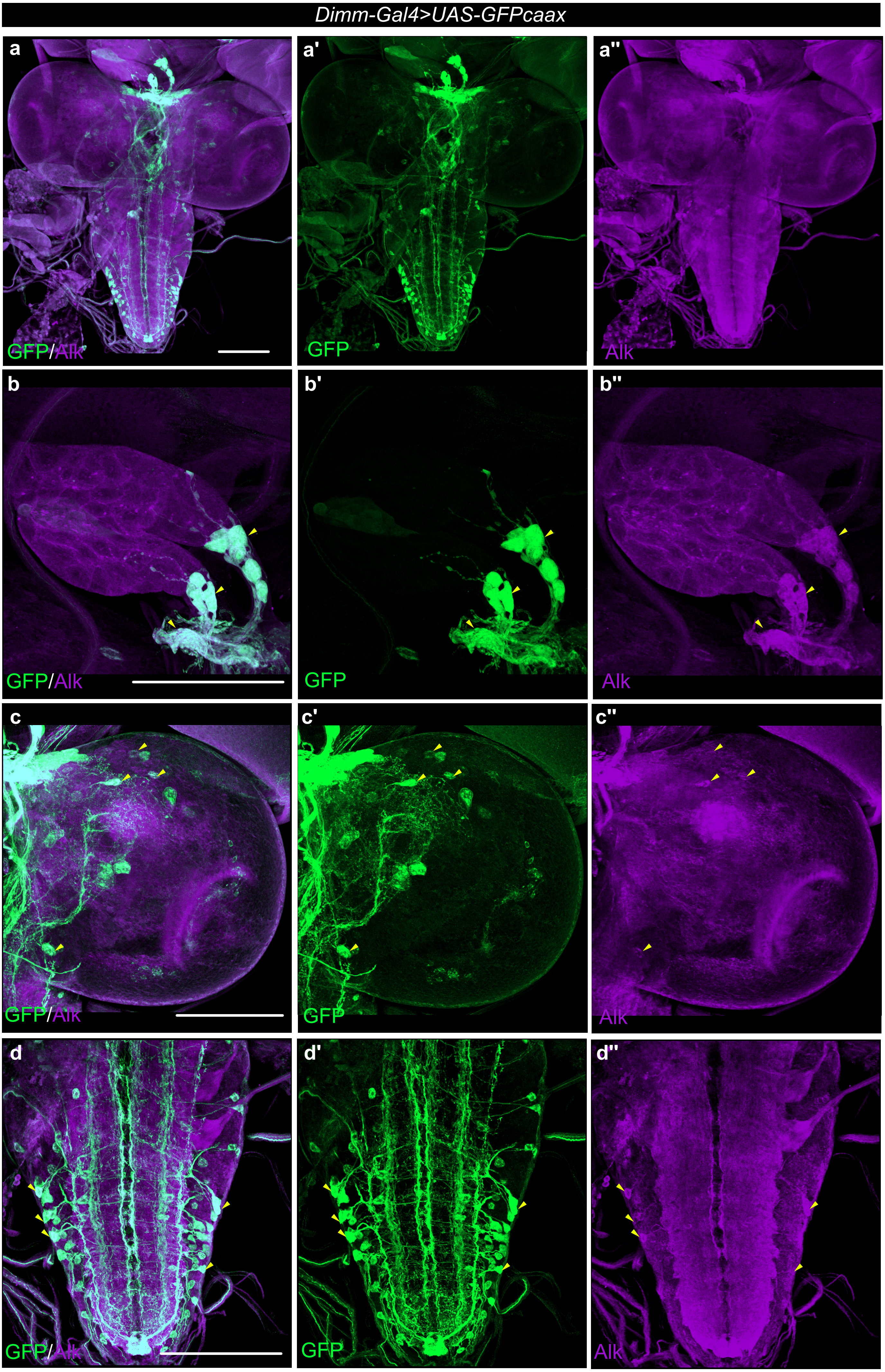

*Sukumar et al., 2024* - **Figure 3 - figure supplement 1**

**a**

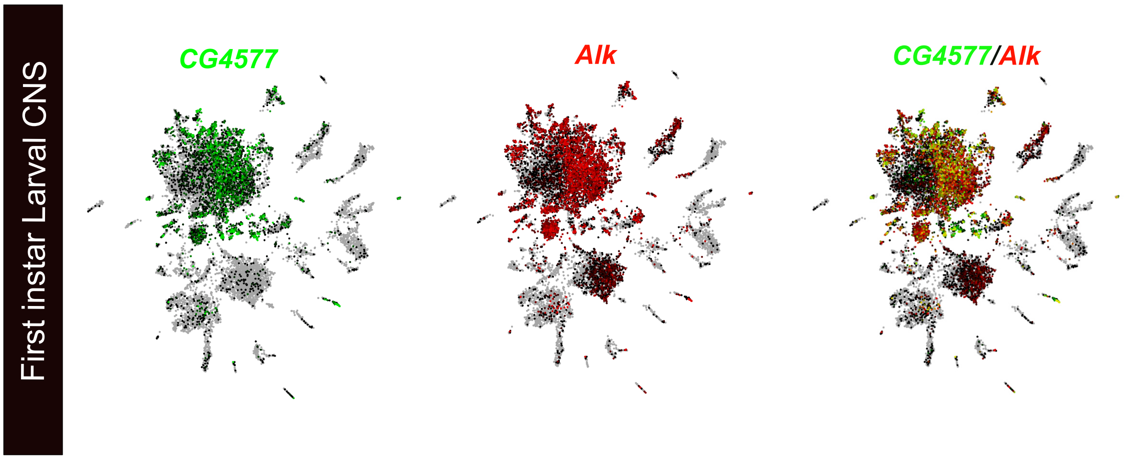


**b**

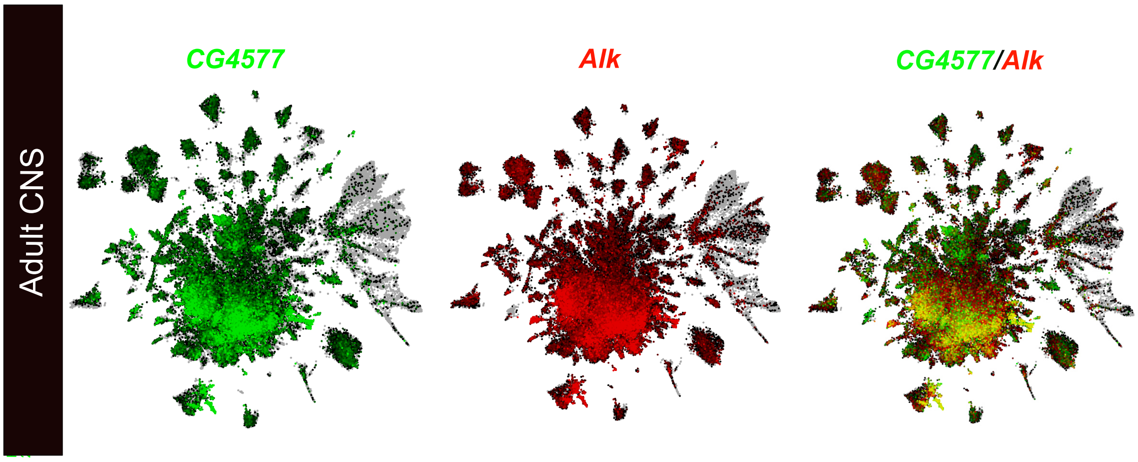


**c**

```
CG4577-PA MPRISLPGSS LLVVLALILC LSLPIRAVPT SVLAQDTELS TSANHKLQQQ QQQQQQHHQQ QQLQQQQQQQ QPATAKRSEE    80
CG4577-PB MPRISLPGSS LLVVLALILC LSLPIRAVPT SVLAQDTELS TSANHKLQQQ QQQQQQHHQQ QQLQQQQQQQ QPATAKRSEE    80
          ********** ********** ********** ********** ********** ********** ********** **********

CG4577-PA ASAVPTADKK SSPEIVPASF SNAPSARNSP VPVSSQQQEQ QQVPSGVYAL PSNEEILAAV AAAAQNSQQQ LQEEPSALEE   160
CG4577-PB ASAVPTADKK SSPEIVPASF SNAPSARNSP VPVSSQQQEQ QQVPSGVYAL PSNEEILAAV AAAAQNSQQQ LQEEPSALEE   160
          ********** ********** ********** ********** ********** ********** ********** **********

CG4577-PA AVASSTMRKR GINYEYNPYS VASSDYGSDV PSGVWSDDYE AAVPVSYGER DLQEIDDYVP ERRVSSSSAR NKAYDNLQNL   240
CG4577-PB AVASSTMRKR GINYEYNPYS VASSDYGSDV PSGVWSDDYE AAVPVSYGER DLQEIDDYVP ERRVSSSSAR NKAYDNLQNL   240
          ********** ********** ********** ********** ********** ********** ********** **********

CG4577-PA LNAEAYLESI PLSVPLTYAN RNYNVDERNK RGIYYNVGTP GGNGASSLGS GSGYNDEGIN LNKYRRFNDM RLKRDTQLNP   320
CG4577-PB LNAEAYLESI PLSVPLTYAN RNYNVDERNK RGIYYNVGTP GGNGASSLGS GSGYNDEGIN LNKYRRFNDM RLKRDTQLNP   320
          ********** ********** ********** ********** ********** ********** ********** **********

CG4577-PA ADMLALVALV EAGERARKES DAESSGPVPM IPSDDLDYAP AGSWFDVPVQ ADYYGAGVPL DNQNQAMPKY EYVPRQHKYS   400
CG4577-PB ADMLALVALV EAGERARKES DAESSGPVPM IPSDDLDYAP AGSWFDVPVQ ADYYGAGVPL DNQNQAMPKY EYVPRQHKYS   400
          ********** ********** ********** ********** ********** ********** ********** **********

CG4577-PA GVNS-RFGSSK QRYMVAKKKR SVNQSQFMNE PVAERGSGYN GEKYF   445
CG4577-PB GVNSPGFGSSK QRYMVAKKKR SVNQSQFMNE PVAERGSGYN GEKYF   446
          ****  ***** ********** ********** ****************
```

*Sukumar et al., 2024* - **Figure 3 - figure supplement 2**

**a**

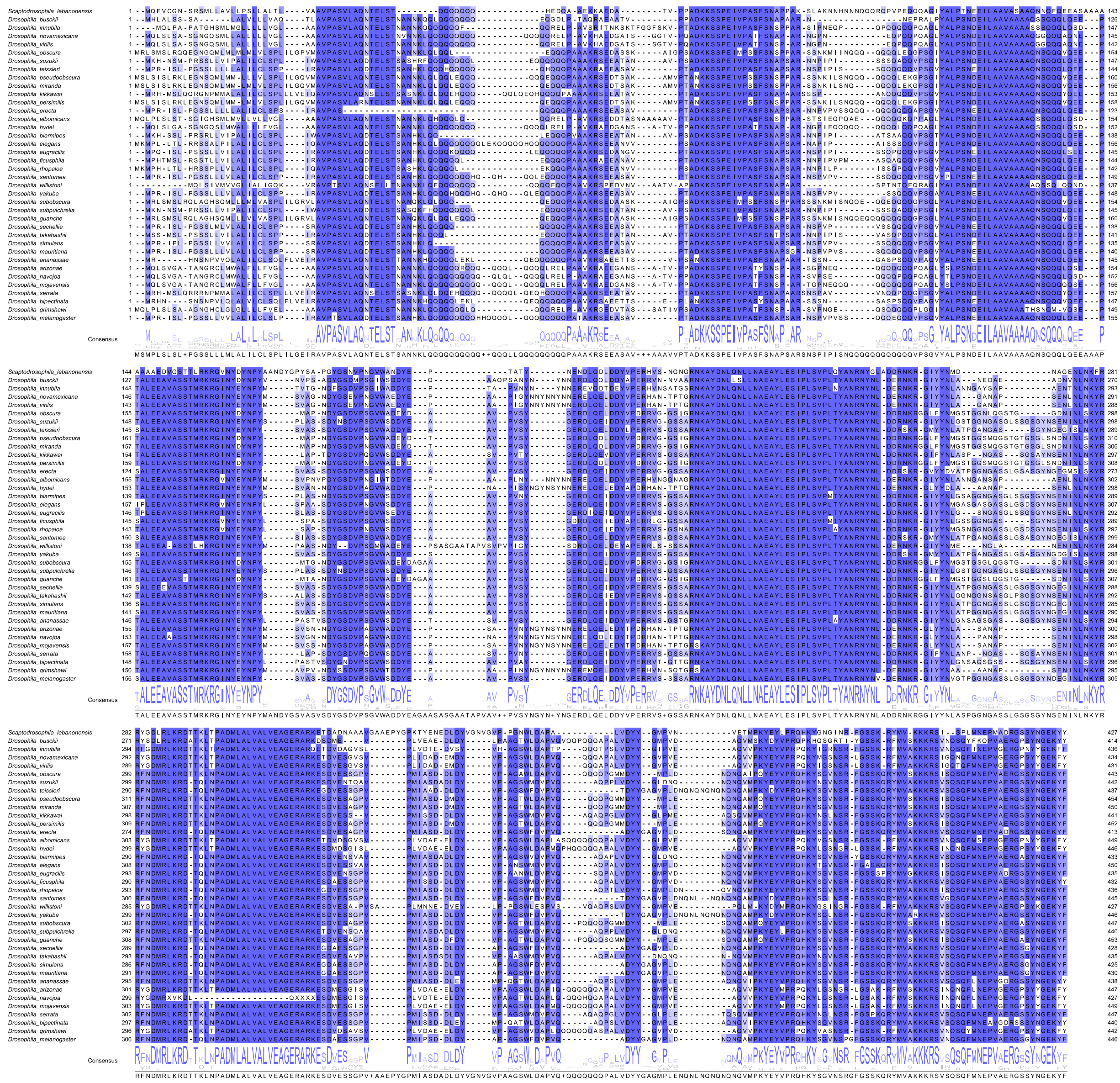

**b**

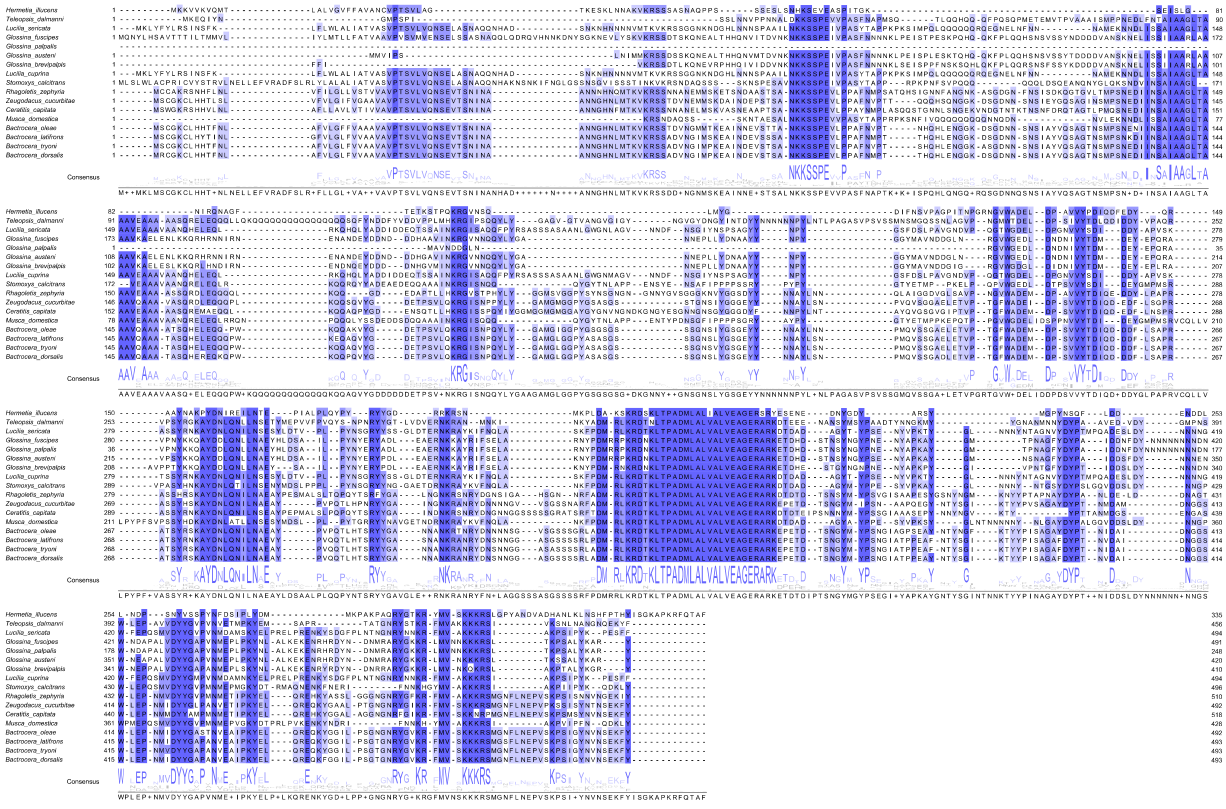

*Sukumar et al., 2024* - **Figure 4 - figure supplement 1**

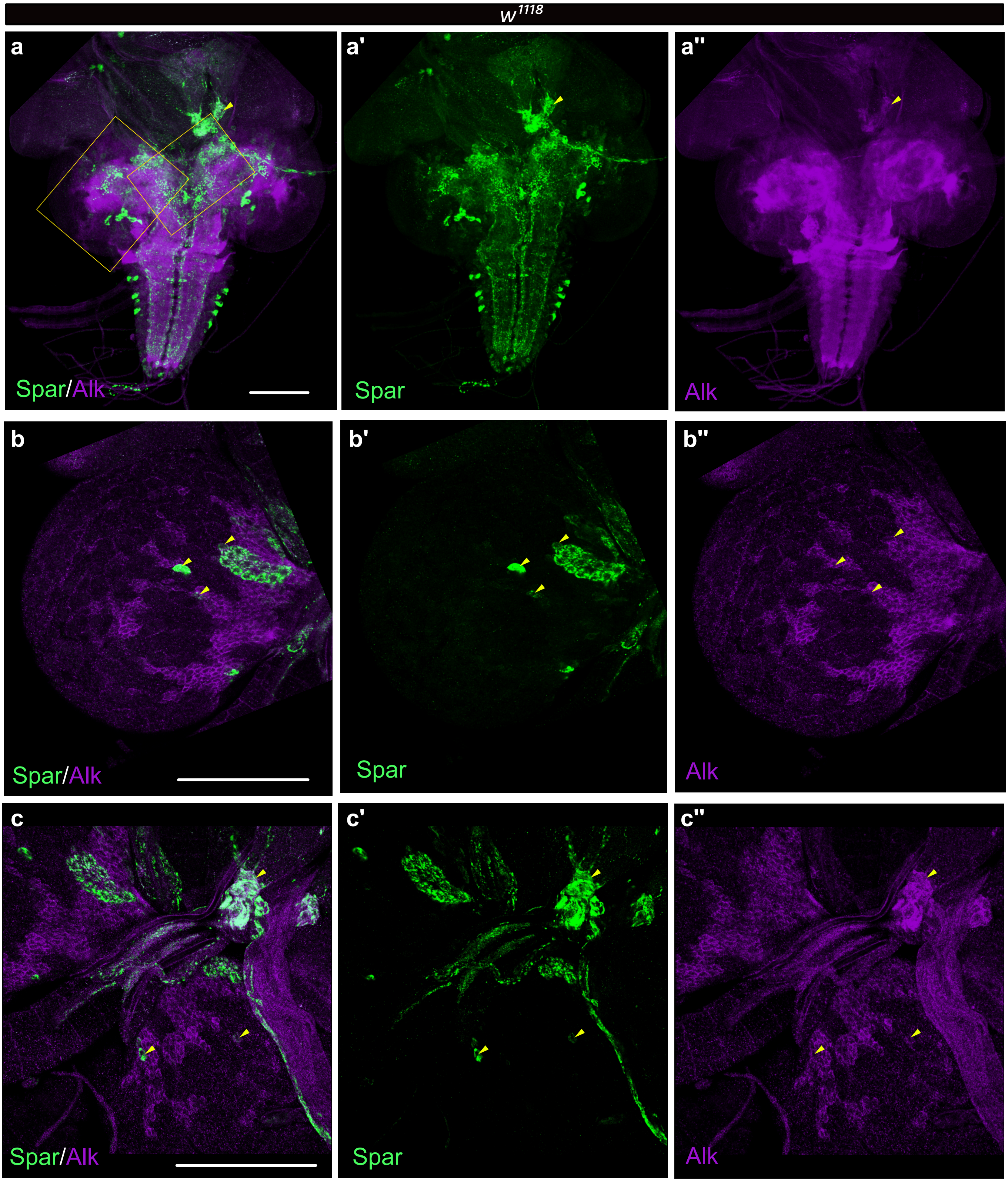

*Sukumar et al., 2024* - **Figure 4 - figure supplement 2**

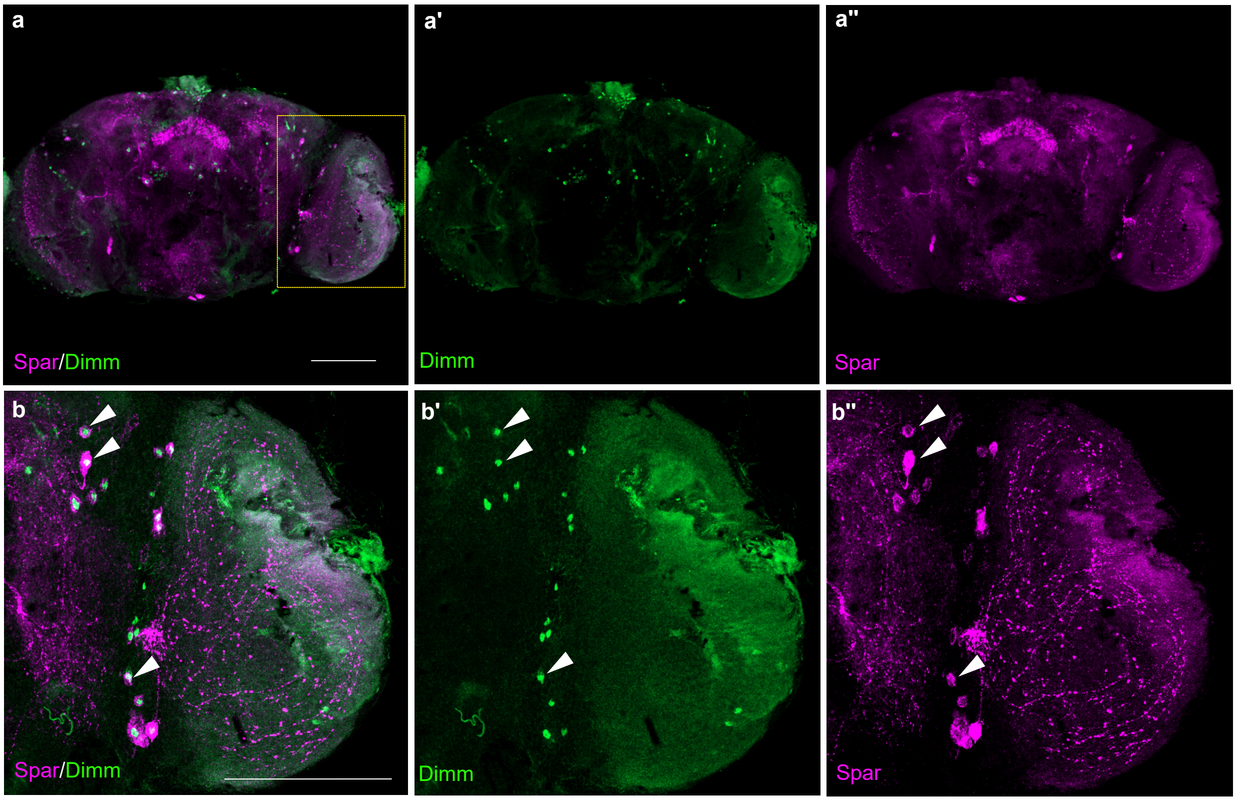

*Sukumar et al., 2024* - **Figure 6 - figure supplement 1**

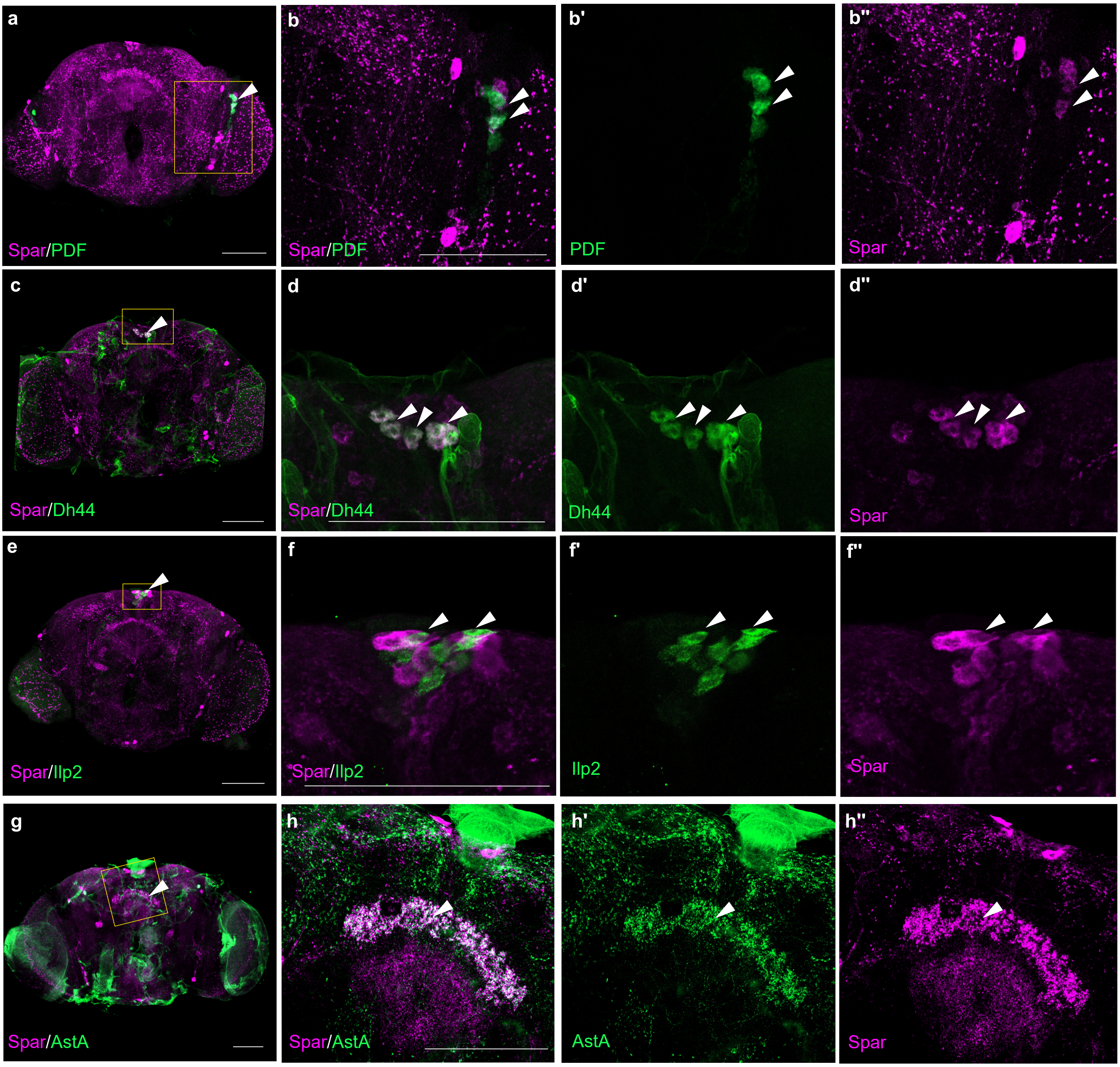

*Sukumar et al., 2024* - **Figure 7 - figure supplement 1**

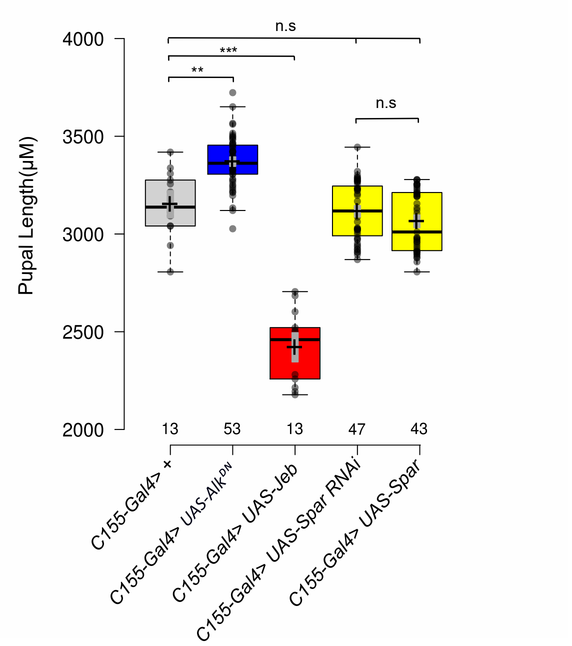

*Sukumar et al., 2024* - **Figure 7 - figure supplement 2**

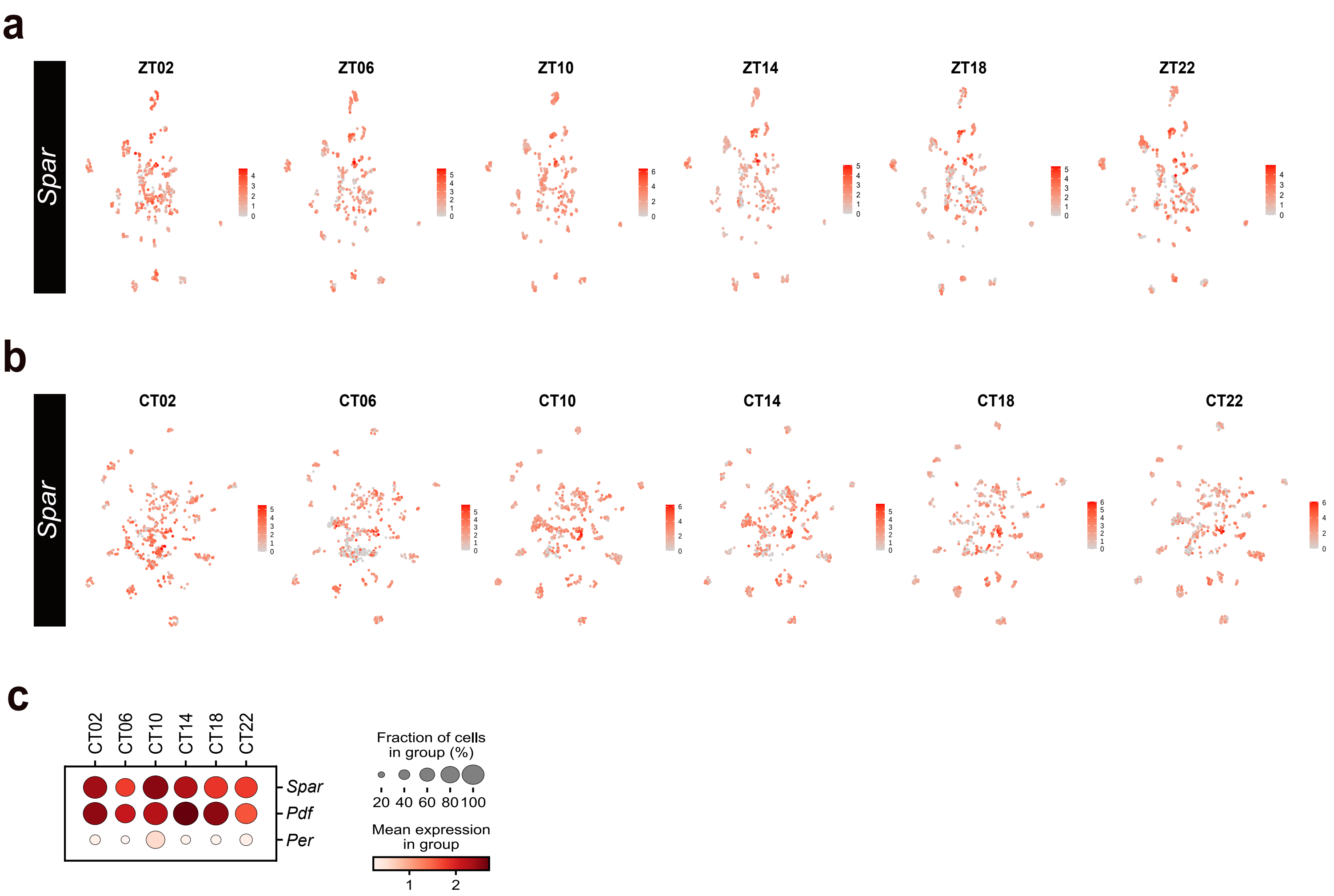

*Sukumar et al., 2024* - **Figure 8 - figure supplement 1**

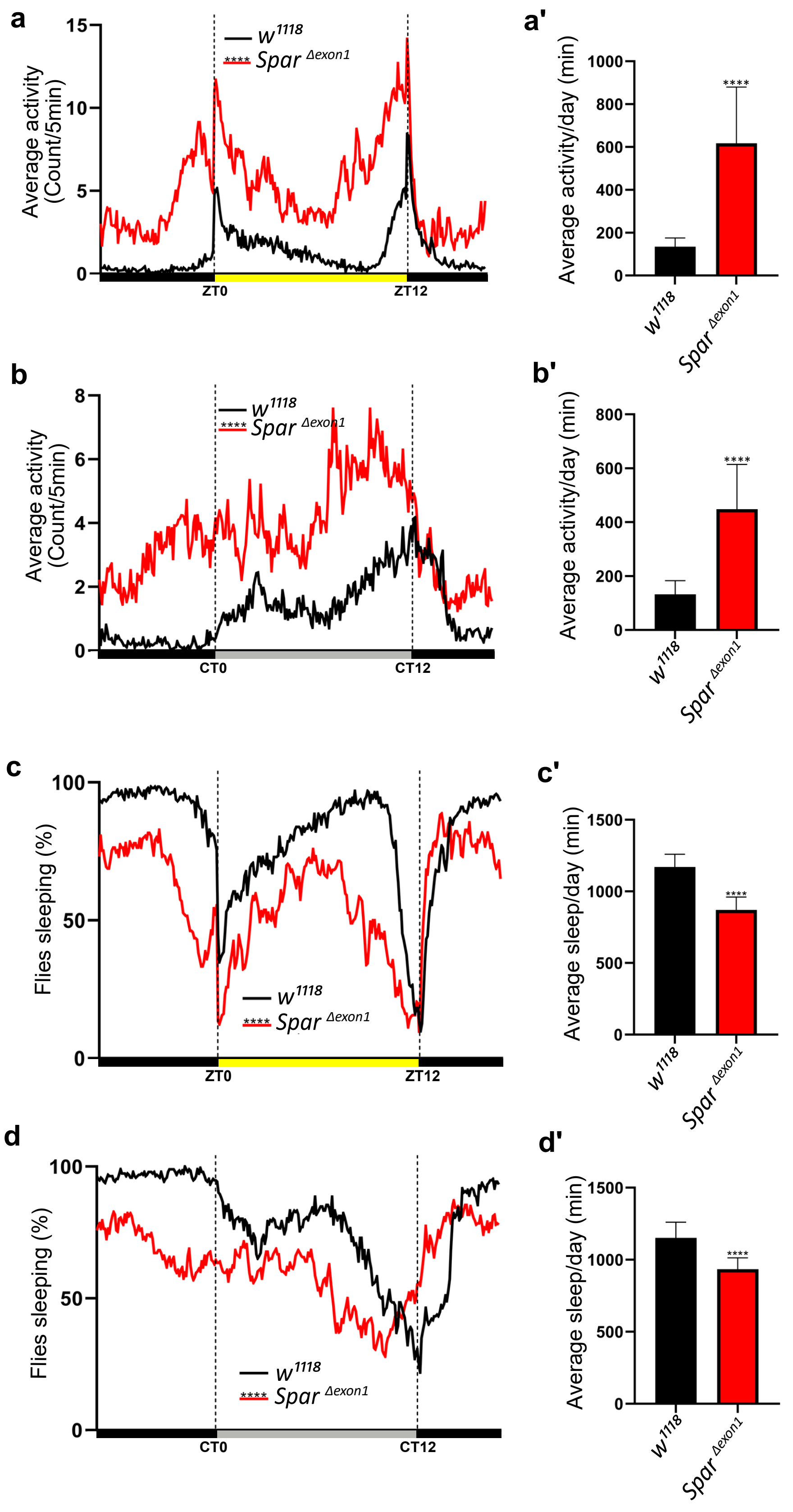

### *Sukumar et al., 2024* - **Figure 8 - figure supplement 2**

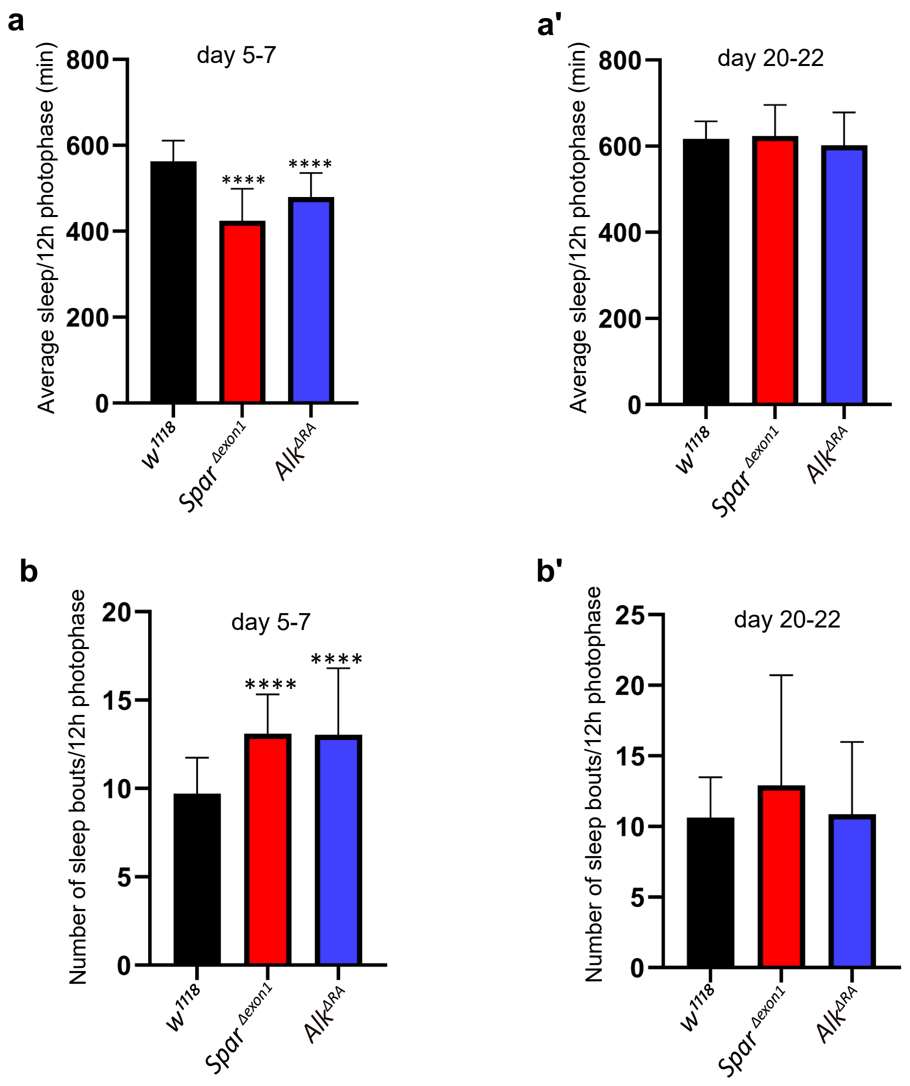

Sukumar et al., 2024 - **Figure 8 - figure supplement 3**

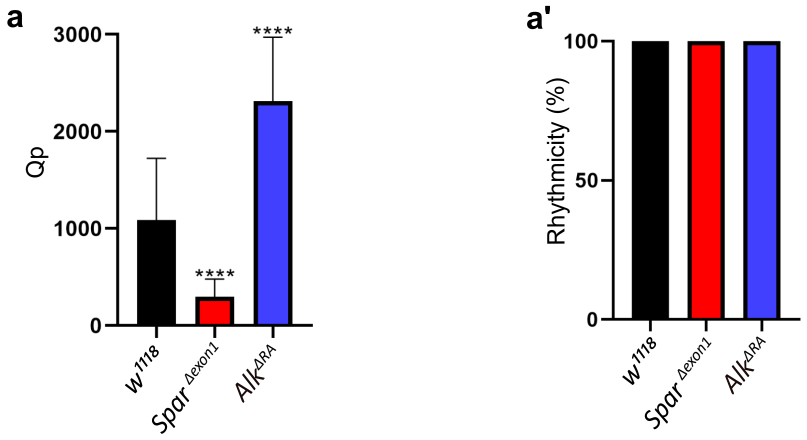

*Sukumar et al., 2024* - **Figure 8 - figure supplement 4**

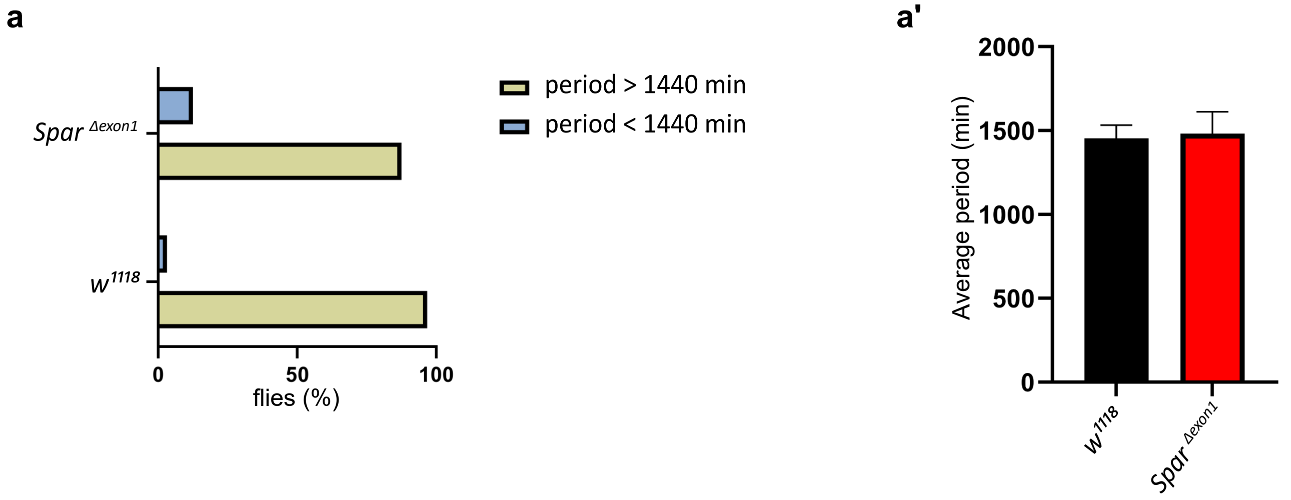

*Sukumar et al., 2024* - **Figure 9 - figure supplement 1**

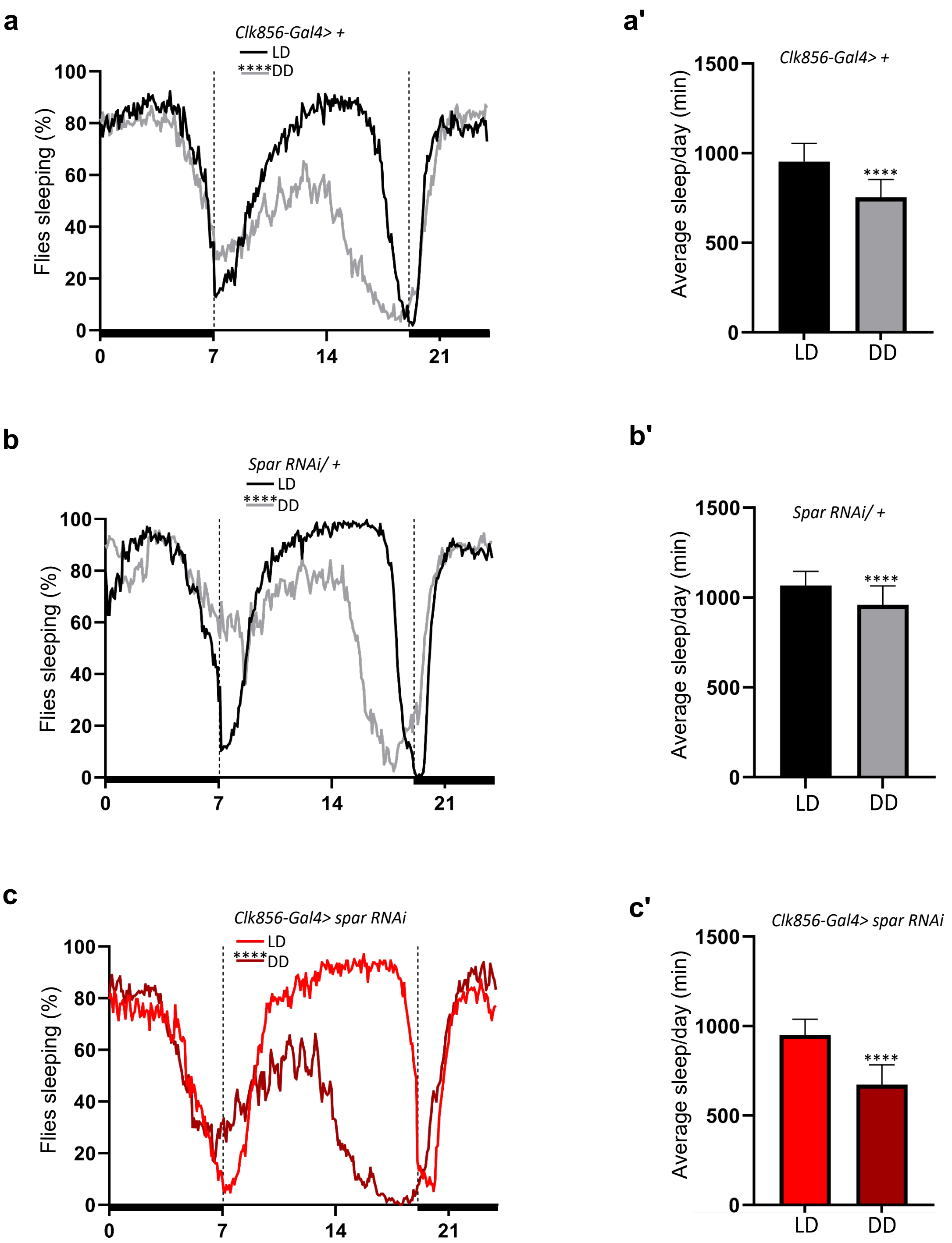

*Sukumar et al., 2024* - **Figure 9 - figure supplement 2**

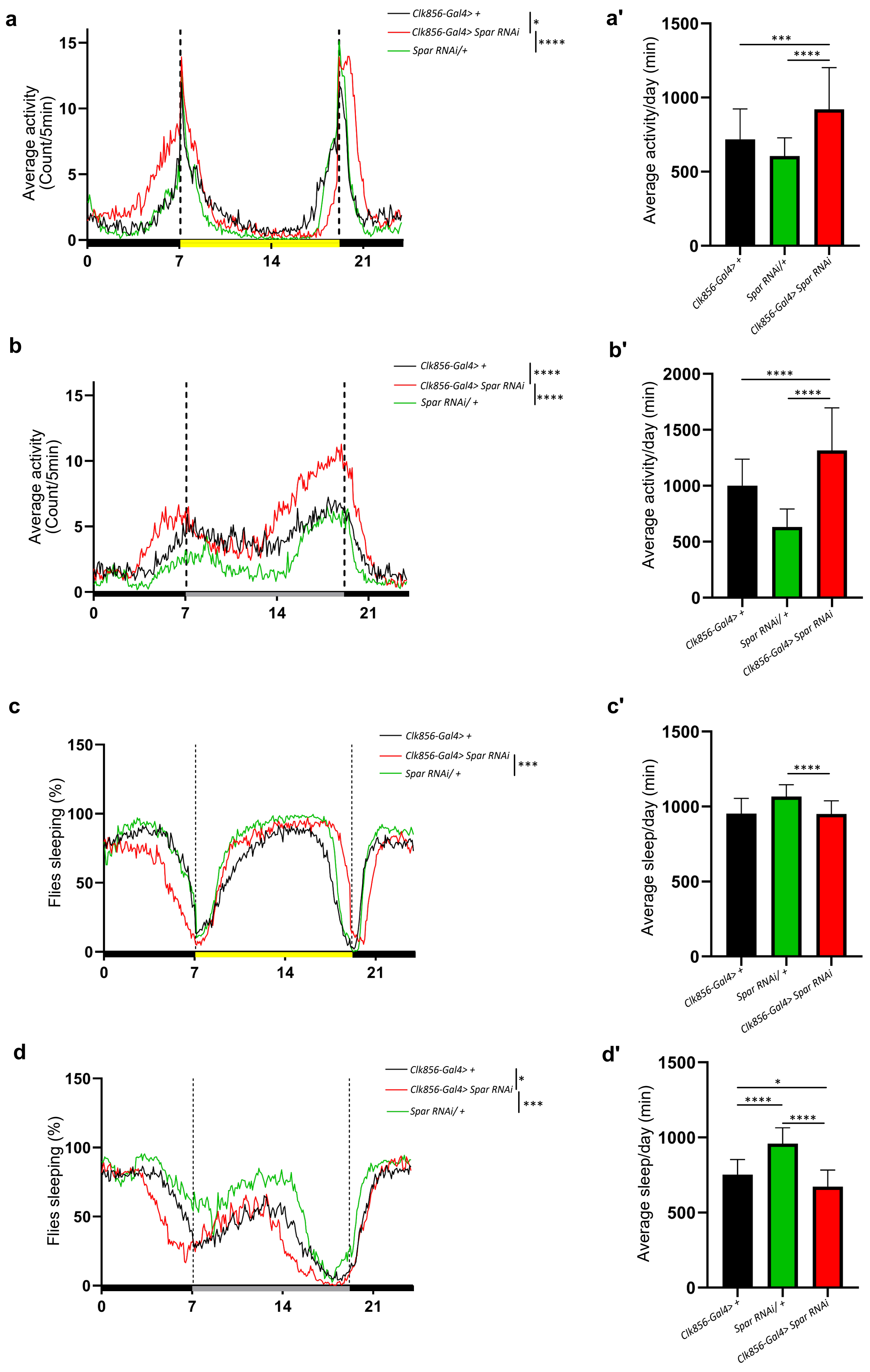

*Sukumar et al., 2024* - **Figure 9 - figure supplement 3**

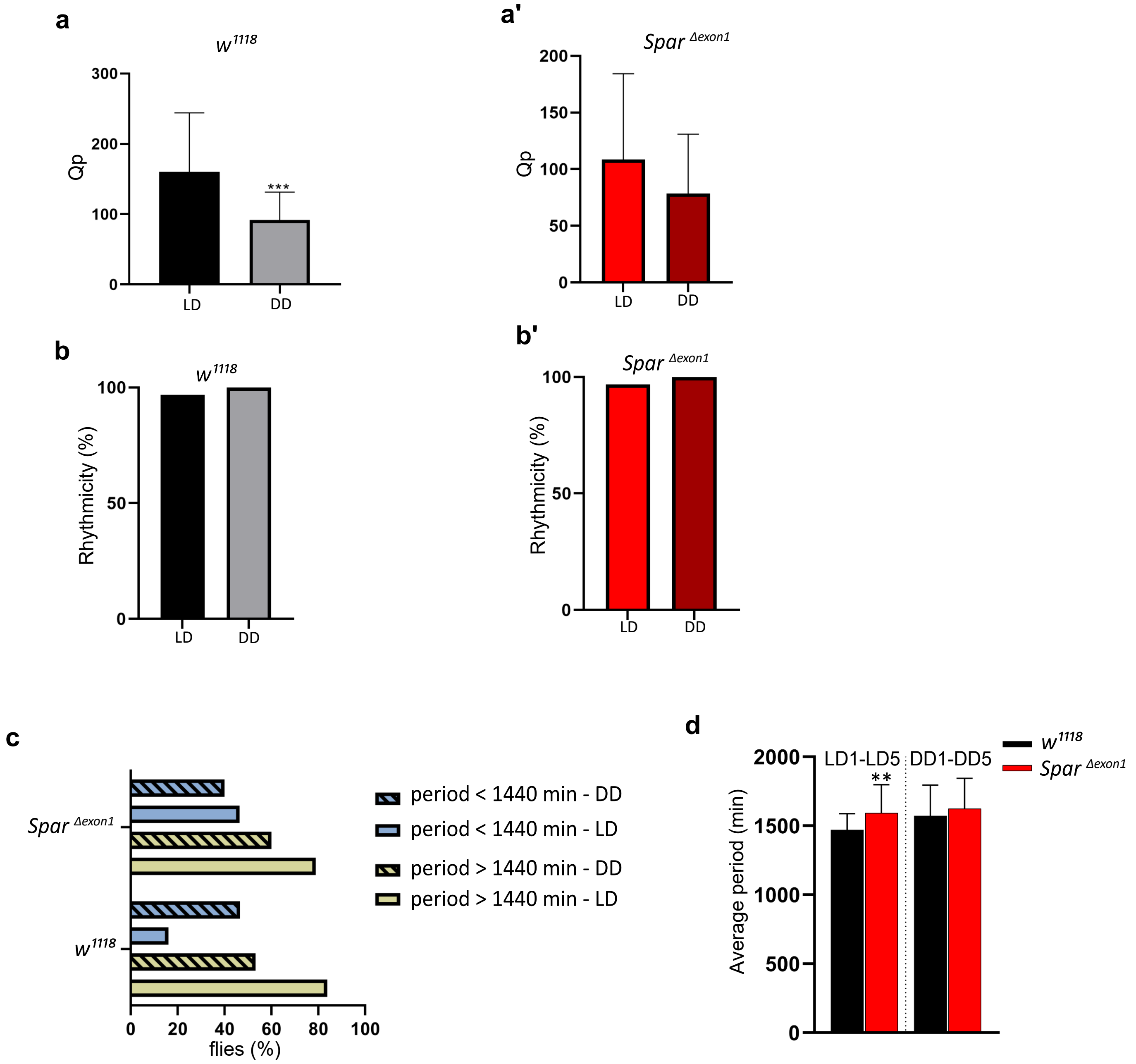

*Sukumar et al., 2024* - **Supplementary Figure 1**

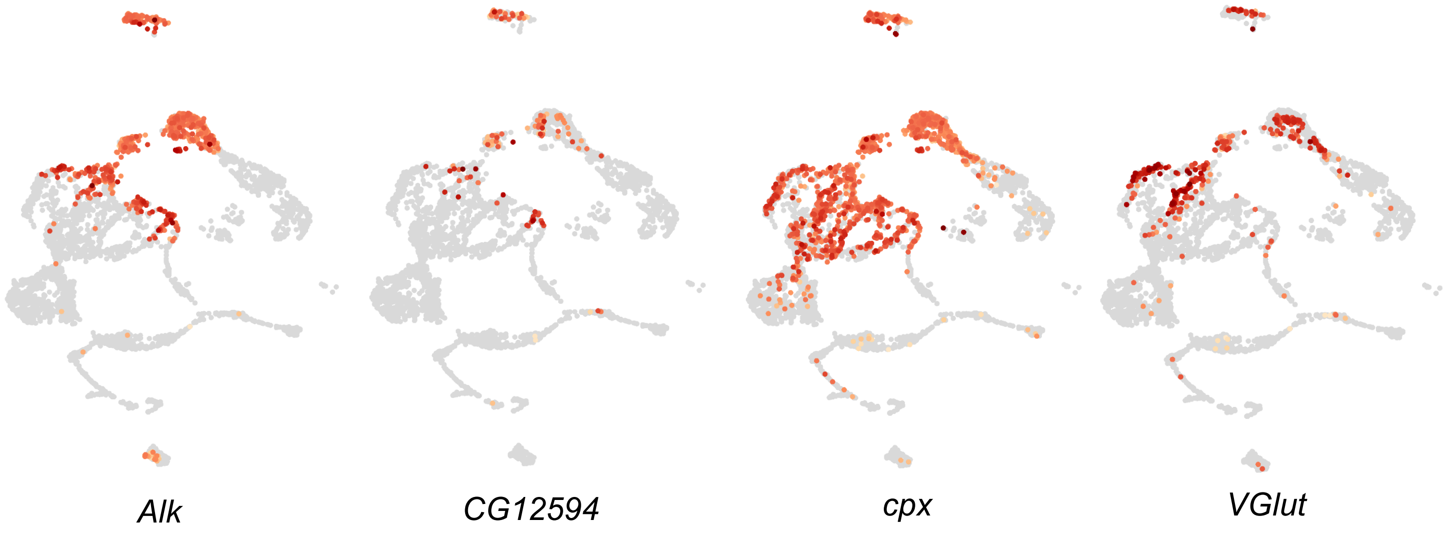
